# Systematics and phylogeography of the Brazilian Atlantic Forest endemic harvestmen *Neosadocus* Mello-Leitão, 1926 (Arachnida: Opiliones: Gonyleptidae)

**DOI:** 10.1101/2021.03.25.436954

**Authors:** Daniel Castro-Pereira, Elen A. Peres, Ricardo Pinto-da-Rocha

## Abstract

*Neosadocus* harvestmen are endemic to the Southern Brazilian Atlantic Forest. Although they are conspicuous and display great morphological variation, their evolutionary history and the biogeographical events underlying their diversification and distribution are still unknown. This contribution about *Neosadocus* includes the following: a taxonomic revision; a molecular phylogenetic analysis using mitochondrial and nuclear markers; an investigation of the genetic structure and species’ diversity in a phylogeographical framework. Our results show that *Neosadocus* is a monophyletic group and comprises four species: *N. bufo*, *N. maximus*, *N. robustus* and *N. misandrus* (which we did not find on fieldwork and only studied the female holotype). There is astonishing male polymorphism in *N. robustus*, mostly related to reproductive strategies. The following synonymies have resulted from this work: *Bunoweyhia variabilis* Mello-Leitão, 1935 = *Neosadocus bufo* (Mello-Leitão, 1926); and *Bunoweyhia minor* Mello-Leitão, 1935 = *Neosadocus maximus* (Giltay, 1928). Most divergences occurred during the Miocene, a geological epoch marked by intense orogenic and climatic events in the Brazilian Atlantic Forest. Intraspecific analyses indicate strong population structure, a pattern congruent with the general behavior and physiological constraints of Neotropical harvestmen.

## Introduction

Harvestmen (Opiliones) constitute one of the largest orders within Arachnida, with more than 6500 species around the world [1]. These organisms exhibit astonishing diversity in the Neotropics (e.g., the family Gonyleptidae, with more than 800 species in over 300 genera [2, 3]). The great majority of harvestmen inhabiting the Brazilian coast are endemic to the Brazilian Rainforest and occupy a limited geographical range [4]. Neotropical harvestmen present low vagility (excepting some species of suborder Eupnoi), are very dependent on high levels of humidity and display limited tolerance to temperature variation [3, 5, 6]. These characteristics have resulted in their generally poor dispersal abilities and narrow species distribution patterns, which reflected in the most detailed subdivision of areas of endemism (AoEs) proposed for the Brazilian Atlantic Forest.(DaSilva et al., 2015 [7]).

*Neosadocus* Mello-Leitão, 1926 (Fig 1) is a gonyleptid genus endemic to the Brazilian Atlantic Forest, known by its conspicuousness and abundance, being the focal taxon of various behavioral studies [8–12].

**Fig 1.**
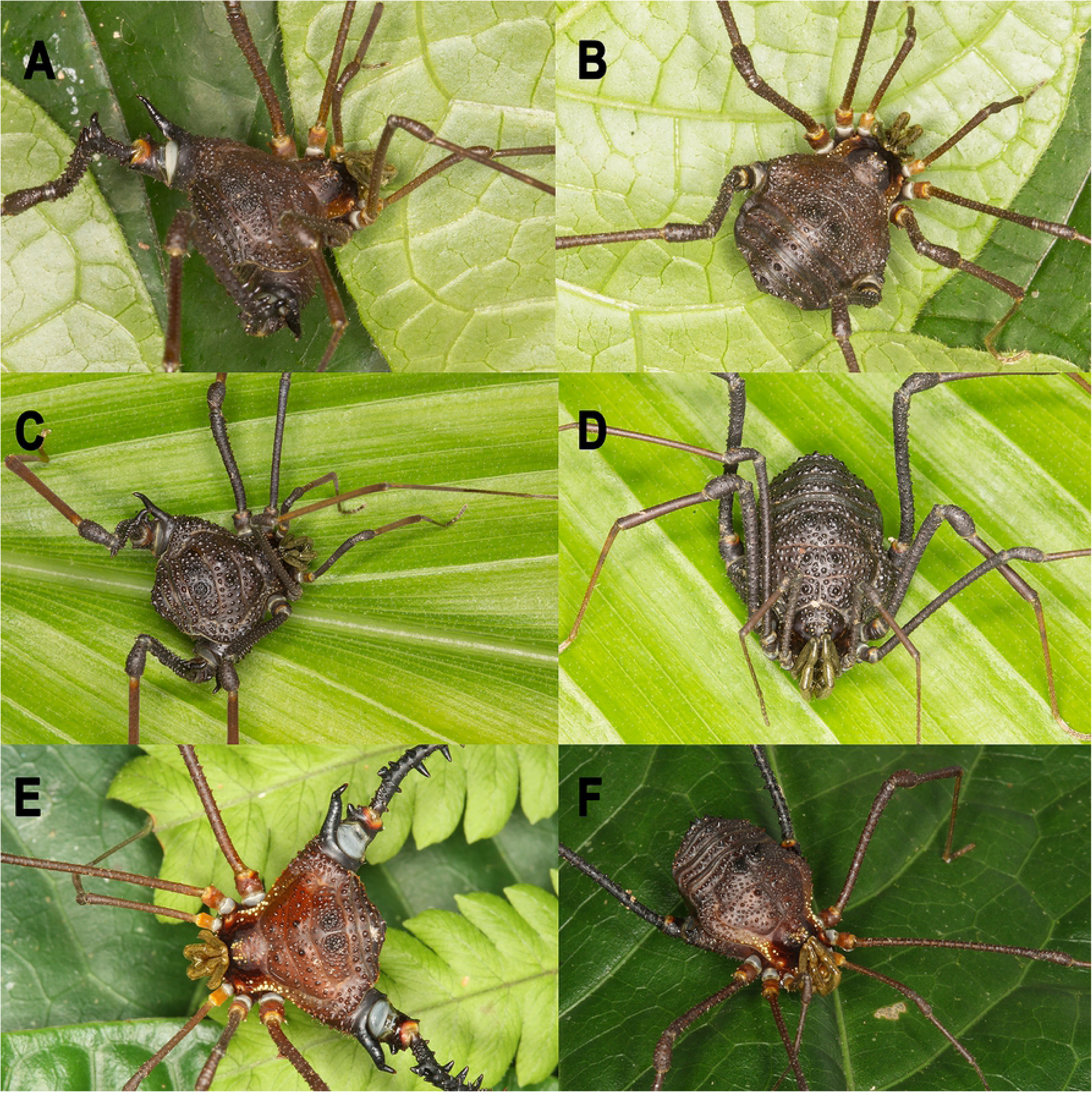
*Neosadocus* living specimens. A, C, E: males; B, D, F: females. A–B. *N. bufo*. C–D. *N. robustus*. E–F. *N. maximus*. All specimens from Brazil, state of São Paulo. *N. bufo* male specimens from Cotia, Portal do Quilombo; *N. robustus* specimens from Ribeirão Grande, Waterfall trail at Maciel; *N. maximus* specimens from Ubatuba, Fazenda Angelim. Photographs A–D by D. Castro-Pereira; photographs E–F by R. Pinto-da-Rocha.

To date, *Neosadocus* comprises four species: *N. bufo* (Mello-Leitão, 1923); *N. maximus* (Giltay, 1928); *N. misandrus* (Mello-Leitão, 1934); and *N. robustus*

(Mello-Leitão, 1936). The type species of the genus is *Sadocus bufo* Mello-Leitão, 1923, transferred to Neosadocus by the same author in 1926 [13]. The history of the genus includes various descriptions of new species and synonymizations. The most recent taxonomic work on *Neosadocus* was performed by Kury in 2003 [2]. The author revalidated *Neosadocus robustus*, previously synonymized with *N. bufo* by Soares in 1944 [14], and transferred *Metagonyleptes misandrus* Mello-Leitão, 1934 to *Neosadocus*, based on the female holotype. Nevertheless, the genus has not been subjected to a taxonomic revision or a phylogenetic study [15] and the validity of the included species is still uncertain due to great variation in morphological characters. Thus, the evolutionary history of the taxon and the biogeographical events underlying its diversification and distribution in the biome are unknown.

*Neosadocus* populations are mostly distributed in humid coastal mountains along Southeastern Brazil (from São Paulo to Paraná states; Fig 2). Known as one of the most diverse and threatened biomes on the planet [16, 17], the Brazilian Atlantic Forest is composed of a mosaic of vegetation physiognomies stretching over extensive latitudinal (3°S to 30°S) and longitudinal (35°W to 60°W) gradients, occupying an area of complex topography and distinct climatic conditions along most of the Brazilian coast [18, 19]. The species richness and endemism levels found in the biome are impressive; among the endemic groups, harvestmen are represented by an exclusive fauna, with about 97.5% of the species endemic to this biome [2, 4, 5]. This biotic and abiotic heterogeneity suggests that multiple historical processes have shaped the regional communities of the Atlantic Forest. Studies on harvestmen have so far attempted to clarify these events by analyzing the relationships among taxa. Using analyses of endemicity and a cladistic biogeography framework, DaSilva et al. (2015, 2017) [7, 20] proposed that possible barriers such as large rivers, mountain ranges and disruptions in forest physiognomy might have influenced the past and present regionalization of the biota. This hypothesis, corroborated by a time-calibrated molecular phylogenetic and phylogeographic data [21–23], indicates that the biological attributes and restricted species’ ranges (restricted to the most humid portions of the Brazilian Atlantic Forest) of Neotropical harvestmen make them suitable for biogeographical investigations [24].

**Fig 2.**
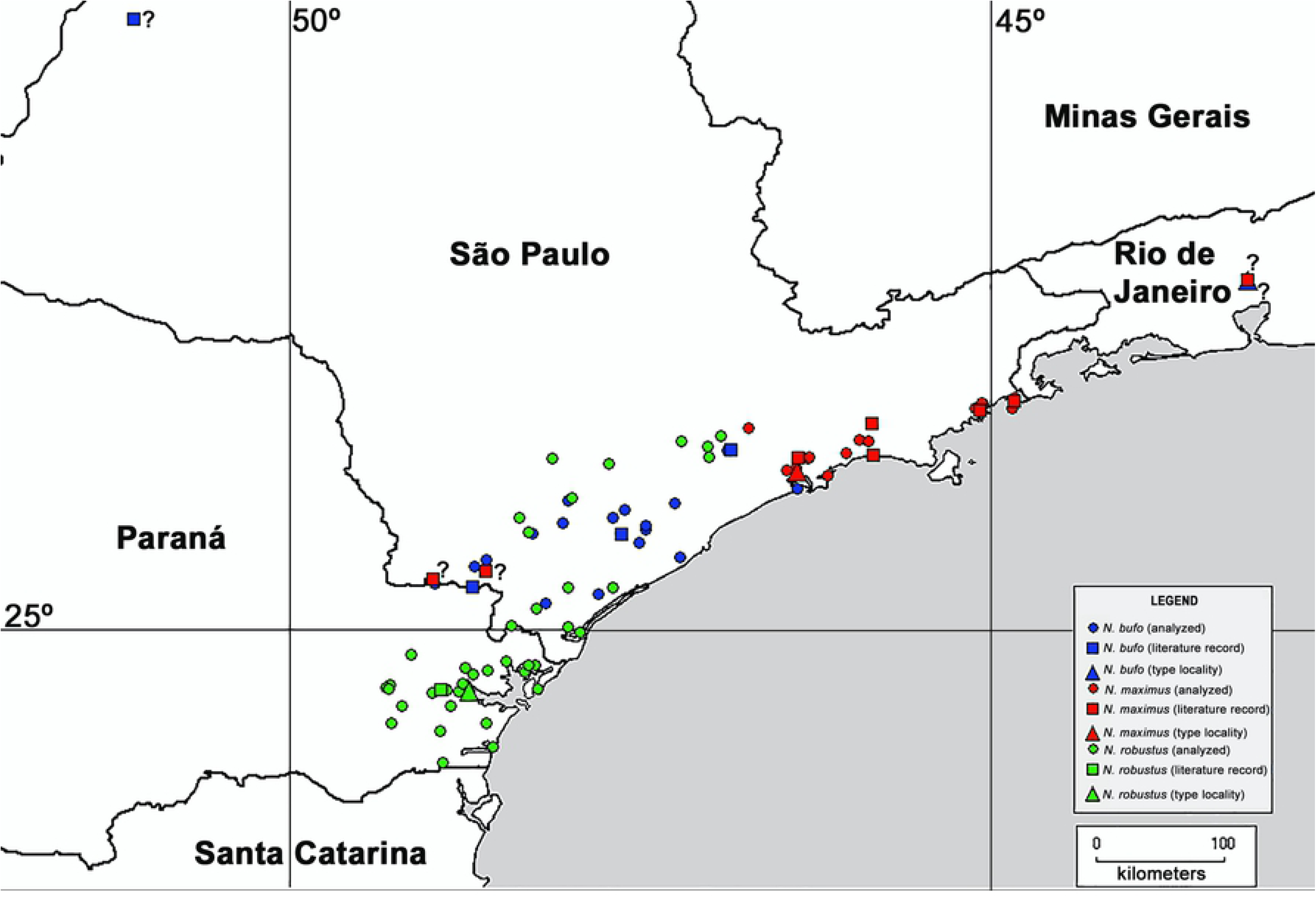
Distribution of *Neosadocus* species. The map indicates localities of specimens collected and used in our morphological and molecular analyses, and locality data of type material and literature records of collection sites. Question marks (?): doubtful records.

In this work, we conducted a taxonomic revision to investigate the validity of *Neosadocus* species using morphological data and inferred their phylogenetic relationships with molecular data. Additionally, we used a phylogeographic framework to infer the possible historical events that promoted the divergences of the major lineages and shaped the population genetic diversity in the genus.

## Materials and methods

### Repositories and taxonomy section

The repositories of the examined material are (within parentheses: Curator; Country, State): IBSP - Instituto Butantan de São Paulo (Dr. Antonio D. Brescovit; Brazil, São Paulo); ISNB - Royal Belgian Institute of Natural Sciences (Dr. Yves Samyn; Belgium, Brussels); MNRJ - Museu Nacional do Rio de Janeiro (Dr. Adriano B. Kury; Brazil, Rio de Janeiro); MZSP - Museu de Zoologia da Universidade de São Paulo (Dr. Ricardo Pinto-da-Rocha; Brazil, São Paulo).

In Taxonomy section, the following abbreviations are implemented: M (male); F (female); DSL (dorsal scutum maximum length); DSW (dorsal scutum maximum width); LDA (length of dorso-basal apophysis of femur IV); LPA (length of paramedian spines of area III).

### Sampling

We analyzed 129 *Neosadocus* specimens collected from 2008 to 2018 in 21 municipalities (32 localities) in the states of São Paulo and Paraná, covering all the known distribution range of the genus (Fig 2, Table 1).

**Table 1.**
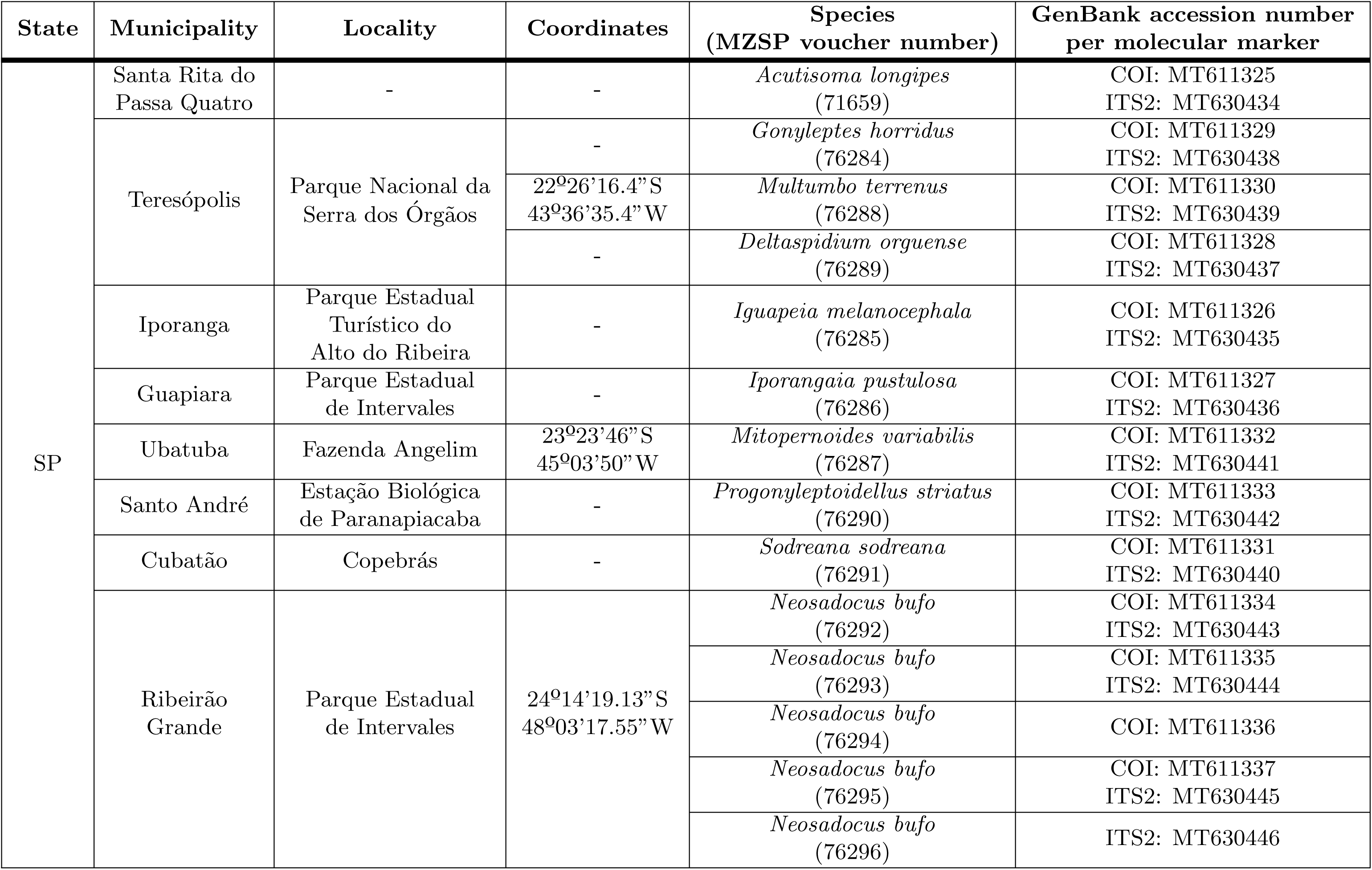
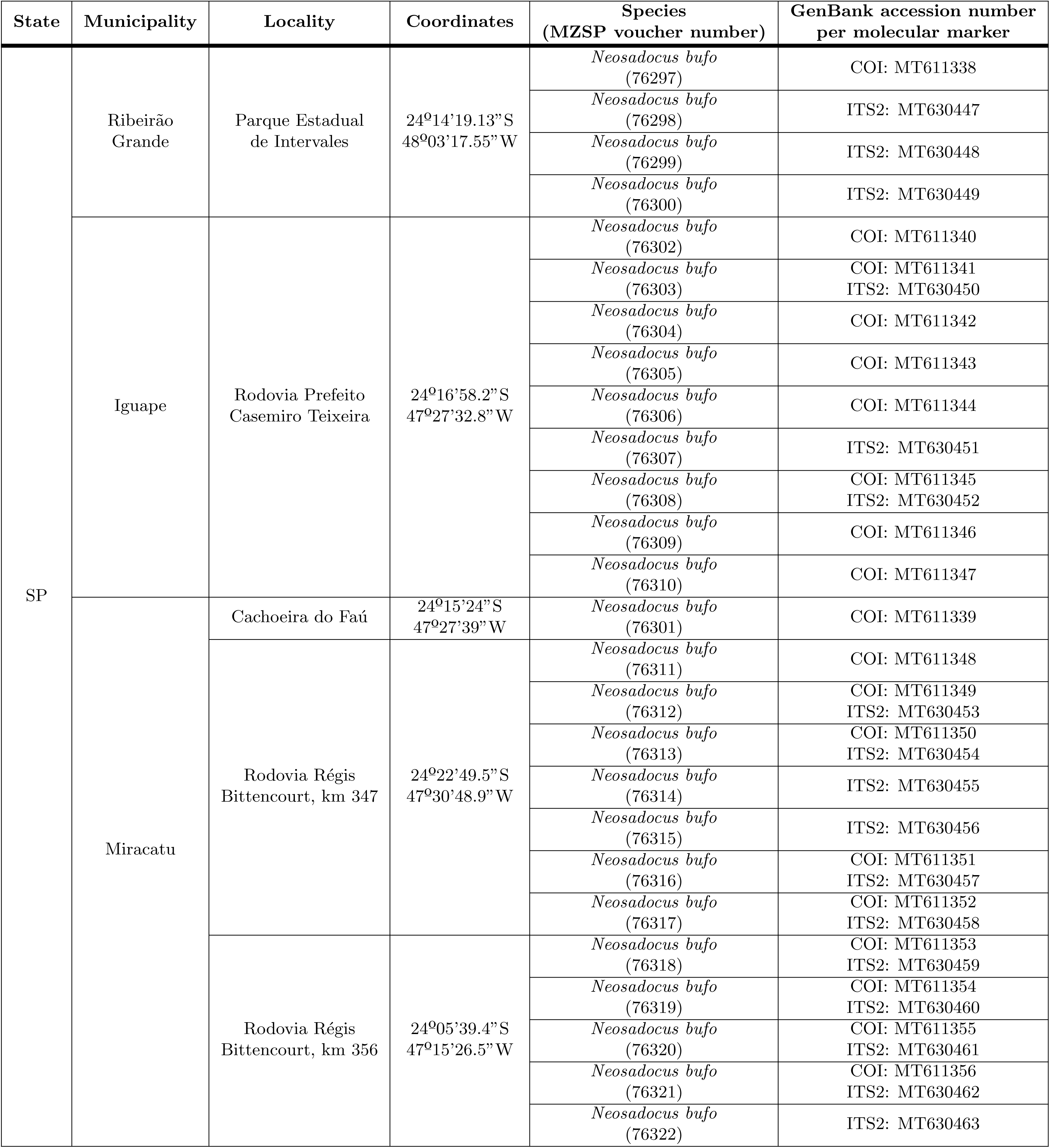
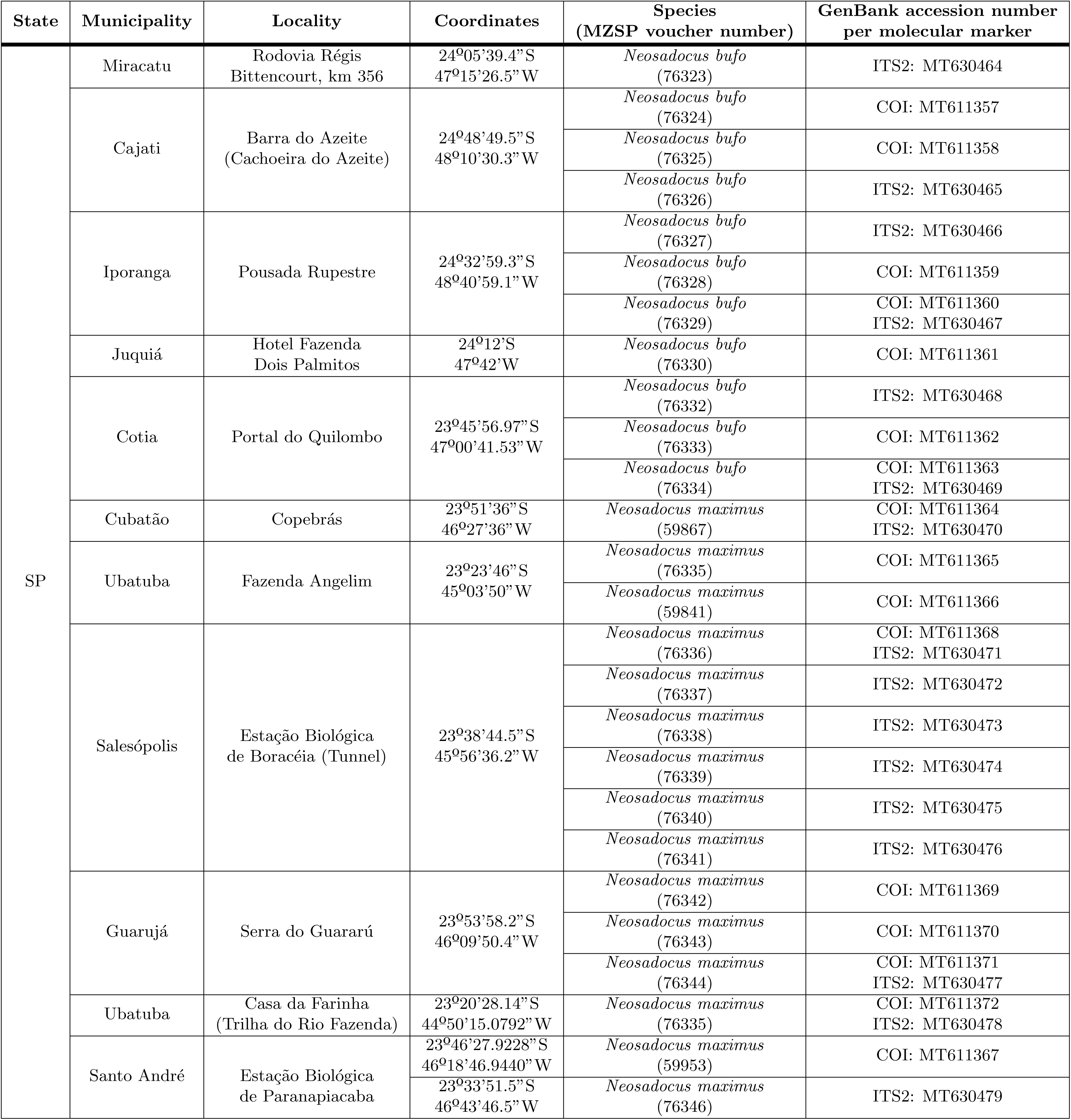
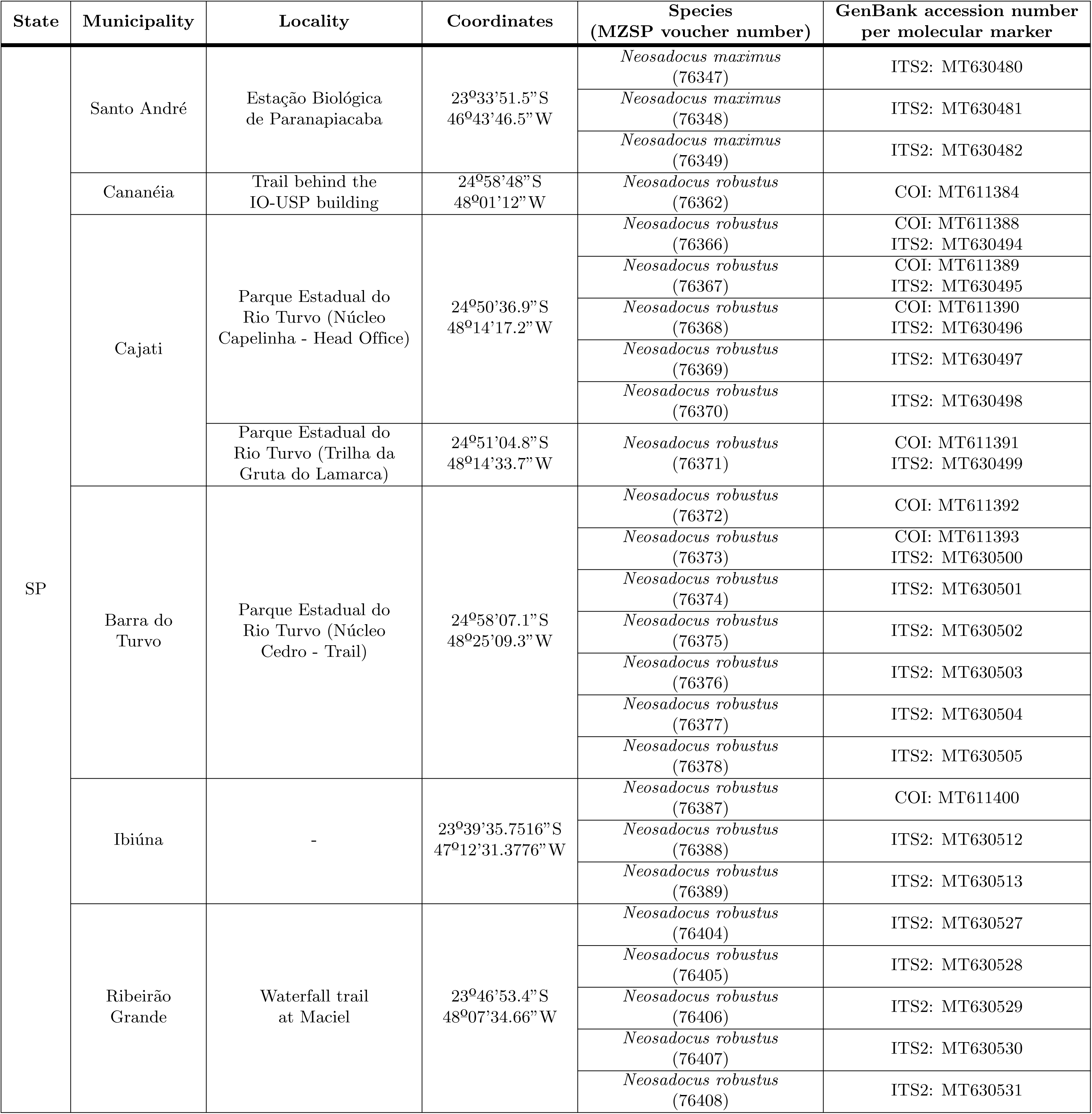
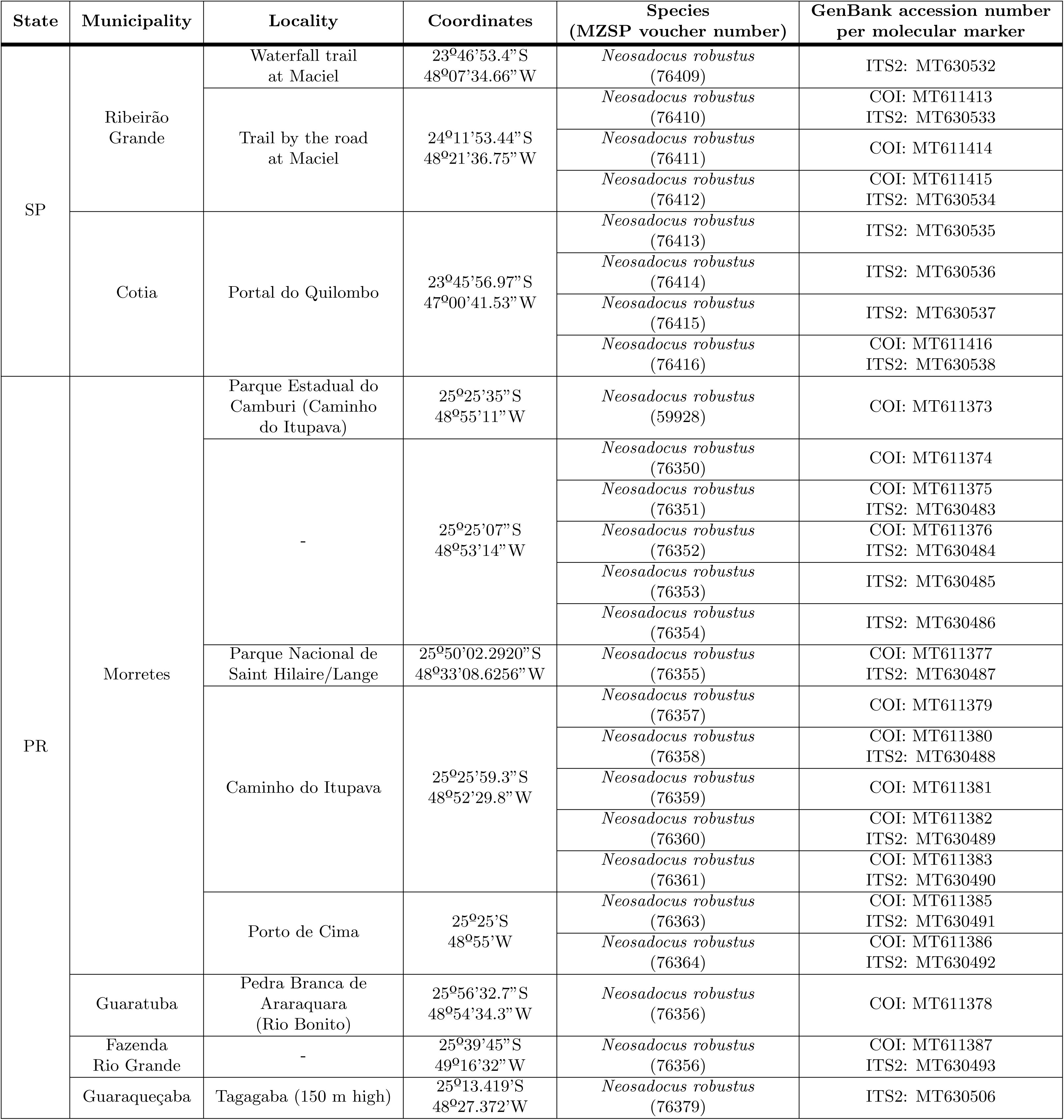
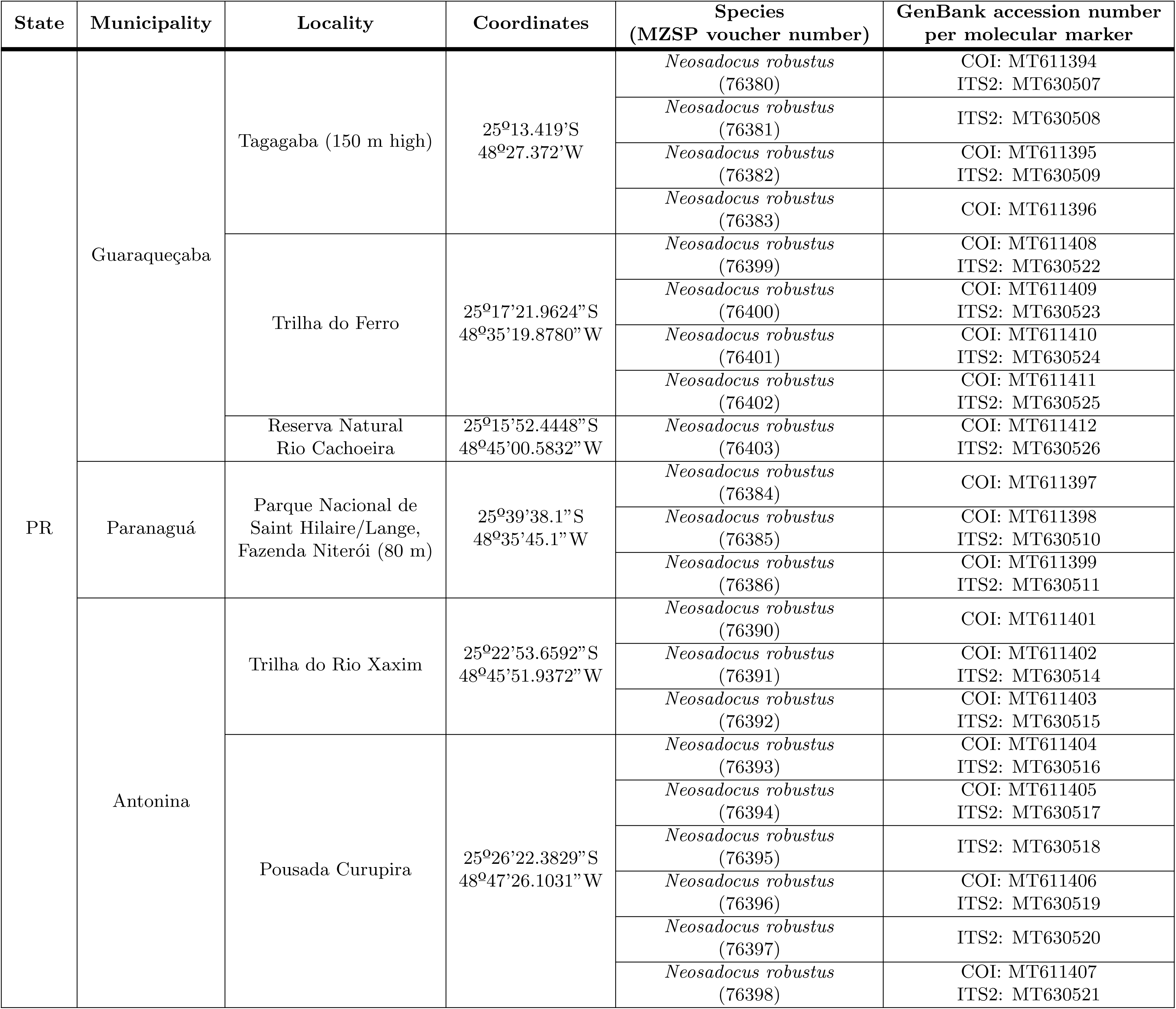
Geographic information, MZSP voucher numbers and GenBank accession numbers of the outgroup species and *Neosadocus* specimens used in this work. Blank spaces (-) indicate missing information. SP = state of São Paulo; PR = Parańa state.

We did not find specimens of *N. misandrus* to study and consequently did not include it in our phylogenetic analysis. The reference to the type locality of the species is vague (Brazil, according to Kury (2003) [2]), which made it impossible to accurately search for it in a specific region.

### Morphological observations and species redescriptions

We analyzed all sequenced male and female specimens of *Neosadocus* and compared them with the type material for species determination under a Leica MS5 stereomicroscope.

The morphological redescriptions were based on the type material of each senior synonym of the valid species. However, we used recently collected specimens to describe the coloration of live animals and the genitalia, and to describe structures that were missing in the type material (see Taxonomy).

All redescriptions follow the same guidelines: the outline of the dorsal scutum is described according to Kury & Medrano (2016) [25]; the topological terms for the appendages are described according to Acosta et al. (2007) [26]; the code for setation of pedipalpal tibia and tarsus are described according to Pinto-da-Rocha (2002) [27]; with respect to tarsi I and II, the numbers within parentheses indicate the number of distitarsi segments; the terminology for the dorsal scutum and leg armature are described according to DaSilva & Gnaspini (2010) [28]; the coloration of 92–96% ethanol-preserved specimens is described according to the NBS/ISCC Color System [http://archive.vn/ufMMn]; the penial macrosetae are described according to Kury & Villarreal M. (2015) [29]; in the descriptions of variation, the numbers within parentheses indicate the number of specimens analyzed; the descriptions of the female only include the characteristics in which they differ from their male counterparts. We followed Pinto-da-Rocha (2002) for the preparation of the male genitalia [27].

The following graphical representation of specimens are provided: photographs of carapace, venter and leg IV of the type material; illustrations of the dorsal scutum, performed using a Leica MZ APO stereomicroscope and a camera lucida; and photographs of the male genitalia, taken with a Zeiss DSM940 scanning electron microscope.

### DNA extraction, amplification, sequencing and alignment

All specimens were stored in ethanol 92–96% at -20°C and the DNA samples were extracted from leg IV muscle tissues using Agencourt DNAdvance kit. We performed polymerase chain reactions (PCRs) for DNA amplification with Thermo Scientific Phusion High-Fidelity DNA Polymerase kit, following manufacturer’s protocols. Two genomic regions were selected for our study: the mitochondrial cytochrome C oxidase subunit I (COI), amplified with the primers dgLCO1490 and dgHCO2198 [30]; and the intronic nuclear internal transcribed spacer 2 (ITS2), with the primers 5.8SF and CAS28Sb1d [31] (S1 Table).

We verified the DNA fragments with 1% agarose gel electrophoresis, purified them with Agencourt AMPure XP kit, and measured DNA concentration of each sample (ng/*µ*L) with a Thermo Scientific NanoDrop 2000 spectrophotometer. For DNA sequencing, we used Applied Biosystems BigDye Terminator v3.1 Cycle Sequencing kit. We also performed DNA precipitation using sodium acetate 3M and ethanol 100% and sent the samples for chromatogram production.

The chromatograms were used for consensus sequences assembly with Phred-Phrap-Consed software pack [32–35]. We manually inspected and edited the sequences in MEGA7 [36] and aligned them with MAFFT software [37], using the default algorithm. Protein-coding COI fragments were examined for stop codons and the sequences of both regions were trimmed to decrease the amount of missing data. Individual ITS2 sequences did not exhibit ambiguous positions (i.e., no heterozygous samples), thus we have not inferred phased nuclear haplotypes. We deposited all sequences in GenBank under accession numbers MT611325 to MT611416 (for COI) and MT630434 to MT630538 (for ITS2; Table 1).

### Phylogenetic analysis and molecular dating

We constructed molecular phylogenetic trees for *Neosadocus* COI and ITS2 sequences using Maximum Likelihood and Bayesian analyses with RAxML [38] and MrBayes (v. 3.2.6) [39], respectively. We first determined the best-fit substitution models for our datasets in jModeltest 2.0 (v. 0.1.1) [40, 41] using the corrected Akaike information criterion (AIC). Nine related Gonyleptidae species (Table 1) were chosen as outgroup, the species *Acutisoma longipes* Roewer, 1913 being the root of the trees. These taxa were chosen according to a previous analysis performed by Pinto-da-Rocha et al. (2014) [42]. In the Maximum Likelihood analyses, we selected each best tree from 100 iterations, and computed the bootstrap values with 1,000 replicates. For the Bayesian inferences, we performed four independent runs with 10,000,000 generations (with chain sampling every 1,000 generations) and determined stationary posterior distribution of parameters by visual inspection in TRACER (v. 1.6). We constructed a 50% majority-rule consensus tree after discarding the first 25% of the trees.

The main lineage divergence events of *Neosadocus* were dated by performing a calibrated multilocus Bayesian analysis using the *BEAST tool [43], available on BEAST software (v. 1.8.0) [44]. We applied the substitution models selected for each dataset and a Yule tree prior under lognormal relaxed (uncorrelated) molecular clock models, since the likelihood ratio tests conducted in MEGA7 rejected the null hypothesis of strict molecular clock for both loci. We used COI and ITS2 mean substitution rates previously estimated for other Neotropical harvestmen [COI: 0.0055/My (0.003–0.008); ITS2: 0.0004/My (0.0002–0.0006); [45]] in normally distributed priors and performed two independent analyses of 100,000,000 generations (with a 10% burn-in). We verified the convergence of runs and checked for parameters’ high effective size values (ESS *>* 200) in TRACER (v. 1.6) and combined the results in LOGCOMBINER (v. 1.8). We annotated both maximum clade credibility (MCC) species and gene trees with TREEANNOTATOR (v. 1.8).

### Population structure and intraspecific analyses

Haplotype networks were constructed for *Neosadocus* COI and ITS2 sequences using the Median-Joining algorithm [46] in PopART software (v. 1.7) [47]. We investigated the genetic structure of *Neosadocus* using Bayesian analyses of population structure (BAPS) in the software BAPS (v. 6.0) [48] and applied the “spatial clustering of groups” option to the COI and ITS2 datasets of each species separately, then inferred the most probable number of clusters (*k*, allowing a range of one to 20 clusters). We performed five replicates for each dataset and the optimal *k* values were based on the highest marginal log-likelihood estimates.

The genetic distances among species and populations (considering each municipality as a population) within each species were calculated in ARLEQUIN (v. 3.5) [49]. To verify the possibility of isolation-by-distance among our samples, we analyzed the correlation between the genetic and geographic distances of the different populations through Mantel tests. We also calculated global *φ*_ST_ per species, pairwise *φ*_ST_ values between populations/BAPS groups, and diversity indices [haplotype (h) and nucleotide (*π*) diversities and number of polymorphic sites] in ARLEQUIN (v. 3.5).

## Results

### Taxonomy

Our morphological observations confirmed the validity of four *Neosadocus* species: *N. bufo*, *N. robustus*, *N. maximus* and *N. misandrus*. This is consistent with the results of the molecular phylogenetic analyses, which recovered the genus as a monophyletic group (the only species absent from the phylogenetic analysis is *N. misandrus*; see Molecular data, phylogenetic analyses and divergence times).

Herein, we present identification keys, diagnoses, and morphological redescriptions for males and females of *N. bufo*, *N. robustus* and *N. maximus*. We provide a complete diagnosis for the female holotype of *N. misandrus*, which is sufficient to distinguish it from the other *Neosadocus* females.

### Gonyleptidae Sundevall, 1833

Gonyleptinae Sundevall, 1833

*Neosadocus* Mello-Leitão, 1833

Figs 1, 3, 4, 5, 6, 7, 8, 9, 10, 11, S1 Fig, S2 Fig

**Fig 3.**
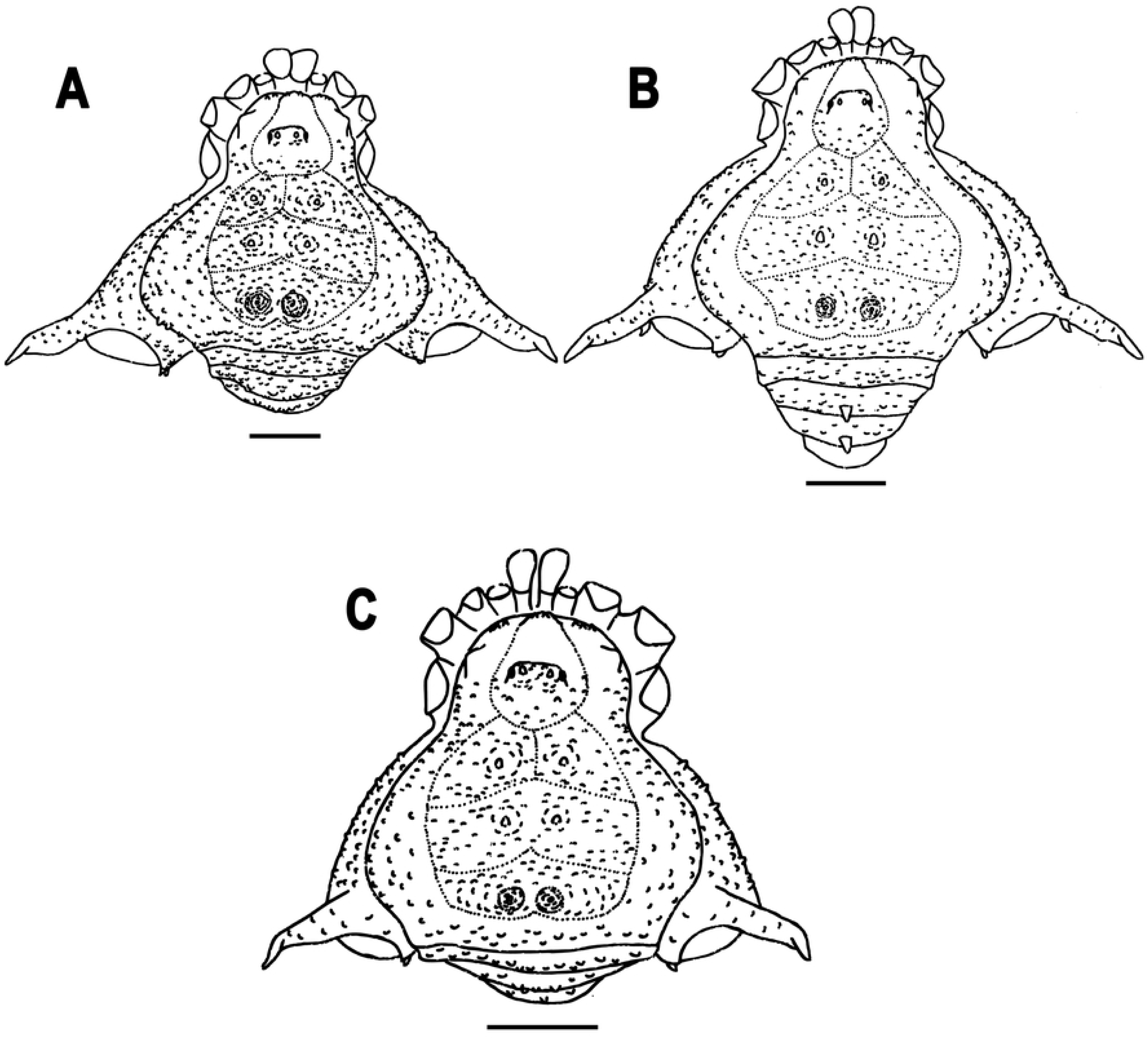
Dorsal habitus of *Neosadocus* males. A. *N. bufo*. B. *N. robustus*. C. *N. maximus*. Scale bars: 3 mm.

**Fig 4.**
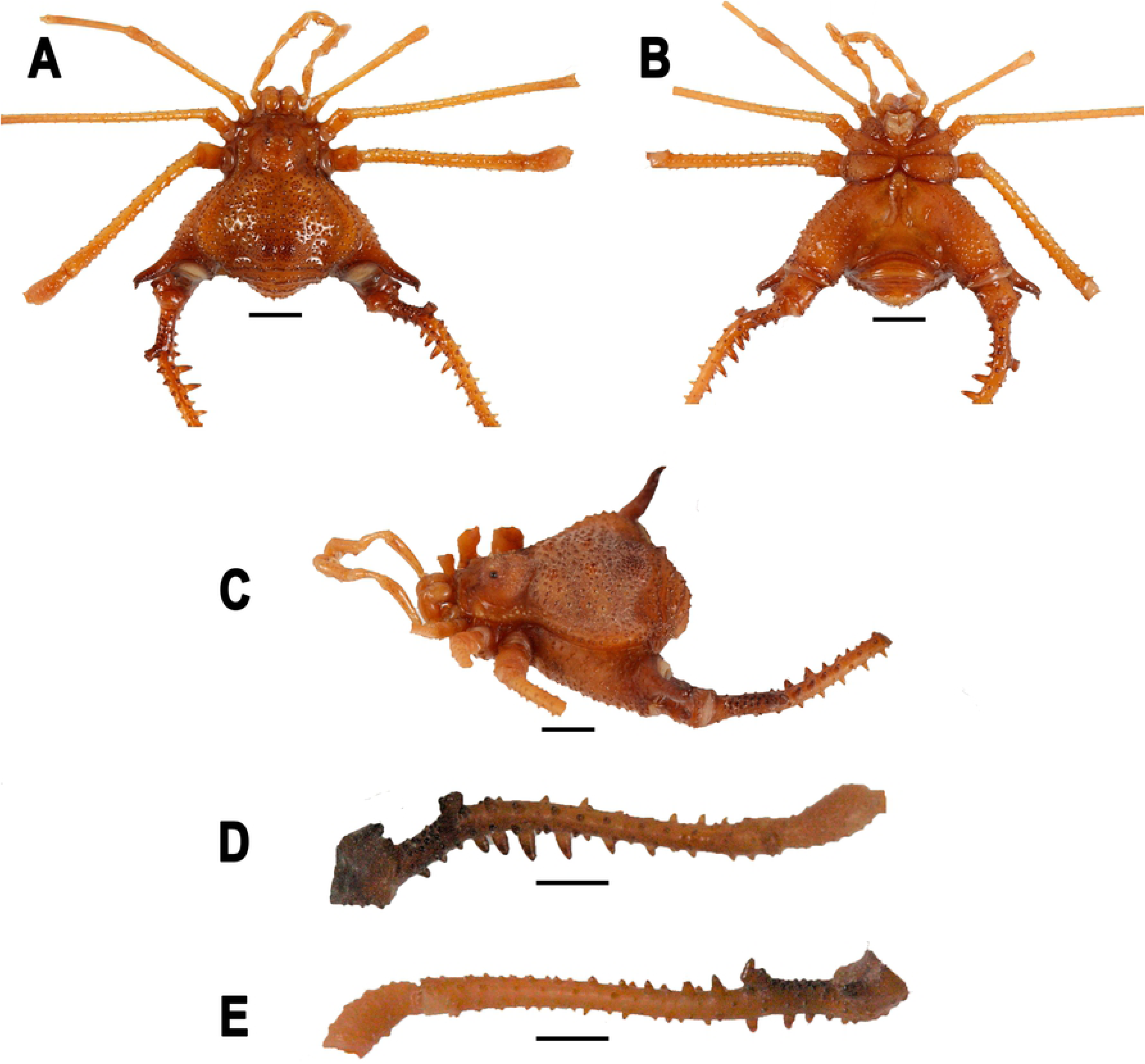
*Neosadocus bufo* (*Bunoweyhia variabilis* male syntype, MNRJ 41803). A. Dorsal view. B. Ventral view. C. Left lateral view. D–E. Right trochanter–patella IV (D: retrodorsal view; E: prolateral view). Scale bar: 3 mm.

**Fig 5.**
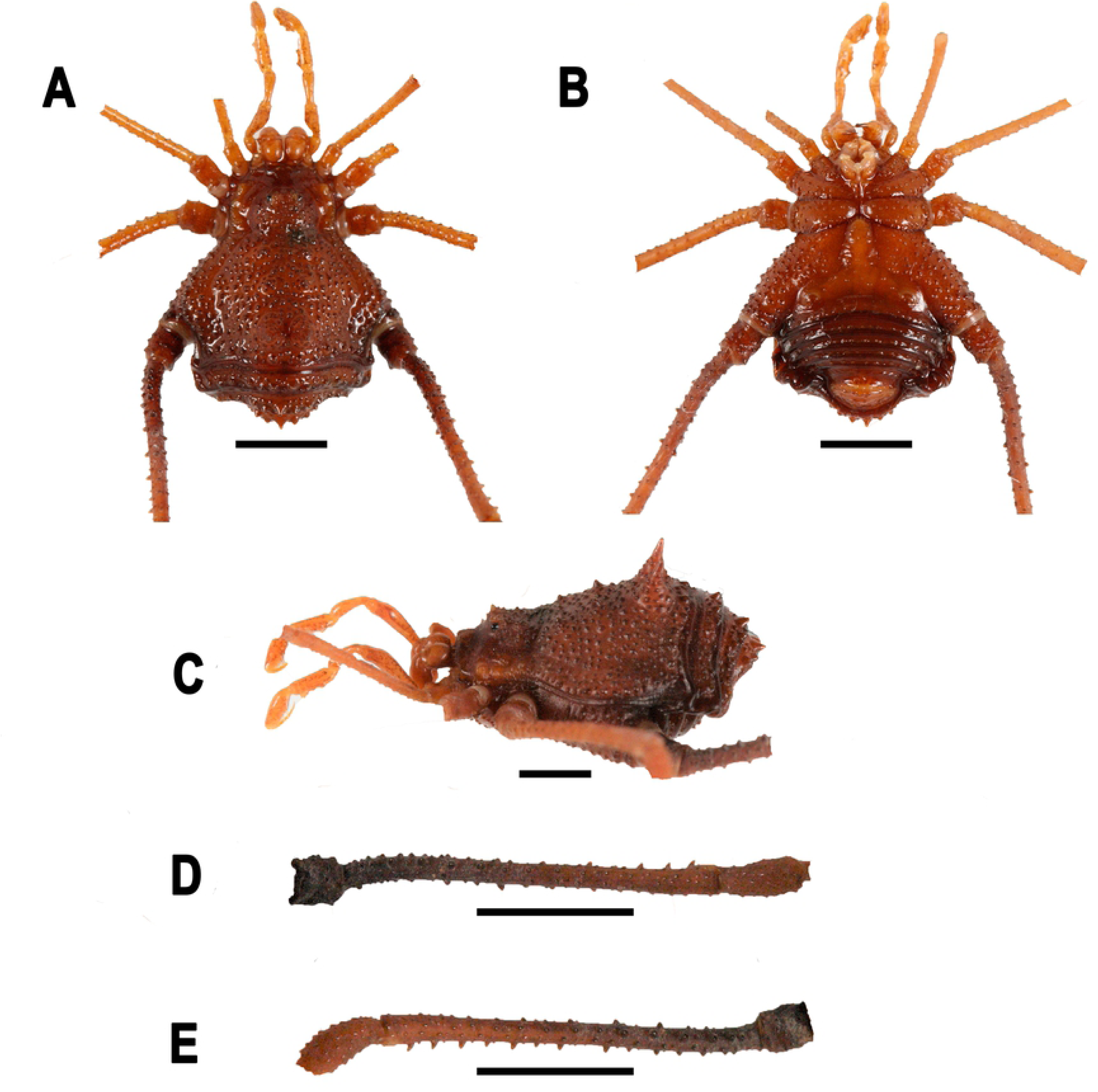
*Neosadocus bufo* (*Bunoweyhia variabilis* female syntype, MNRJ 41803). A. Dorsal view. B. Ventral view. C. Left lateral view. D–E. Right trochanter–patella IV (D: retrolateral view; E: prolateral view). Scale bars: 3 mm.

**Fig 6.**
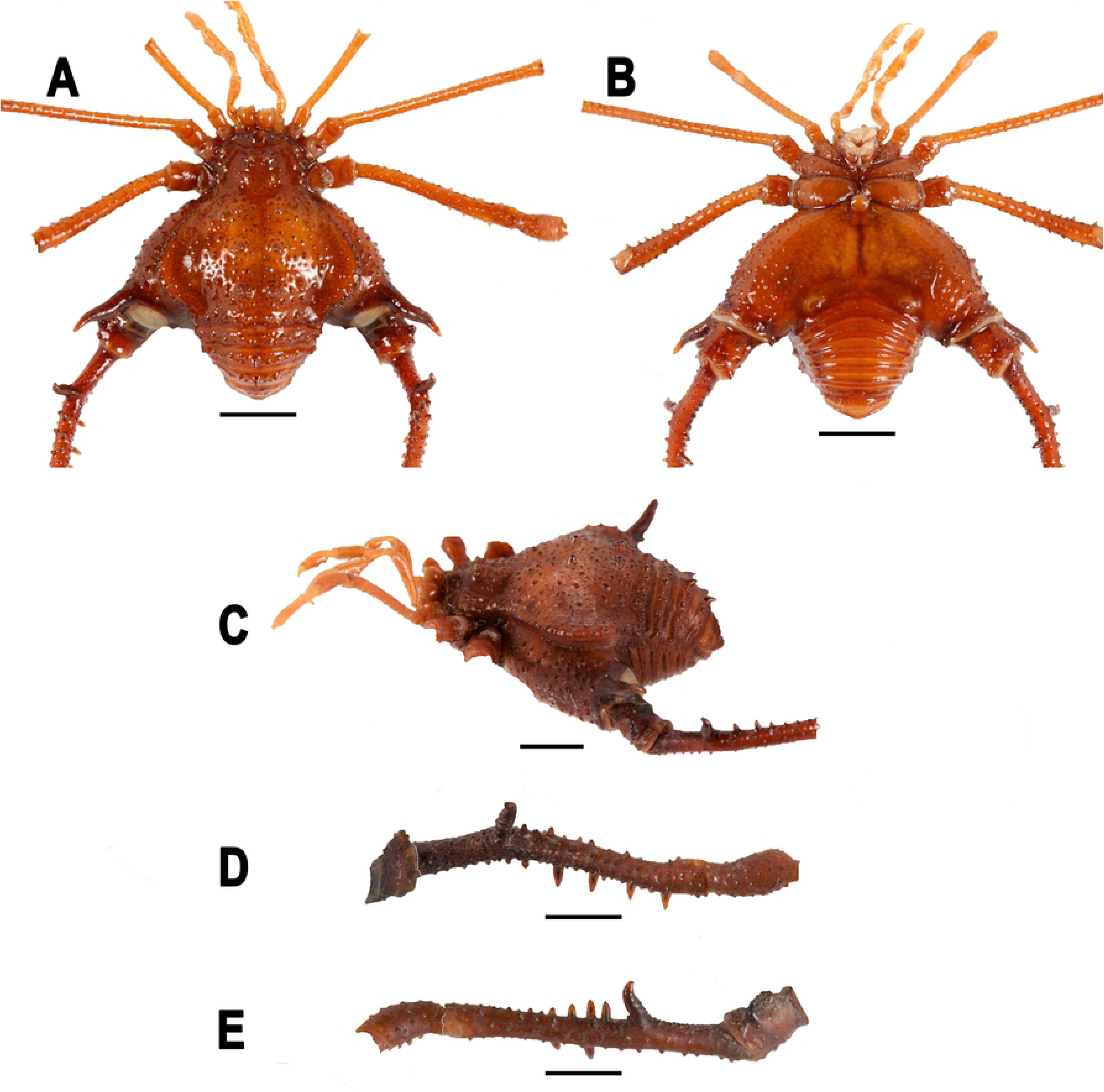
*Neosadocus robustus* (*Ilhania robusta* male lectotype, MNRJ 42289). A. Dorsal view. B. Ventral view. C. Left lateral view. D–E. Right trochanter–patella IV (D: retrolateral view; E: prolateral view). Scale bars: 3 mm.

**Fig 7.**
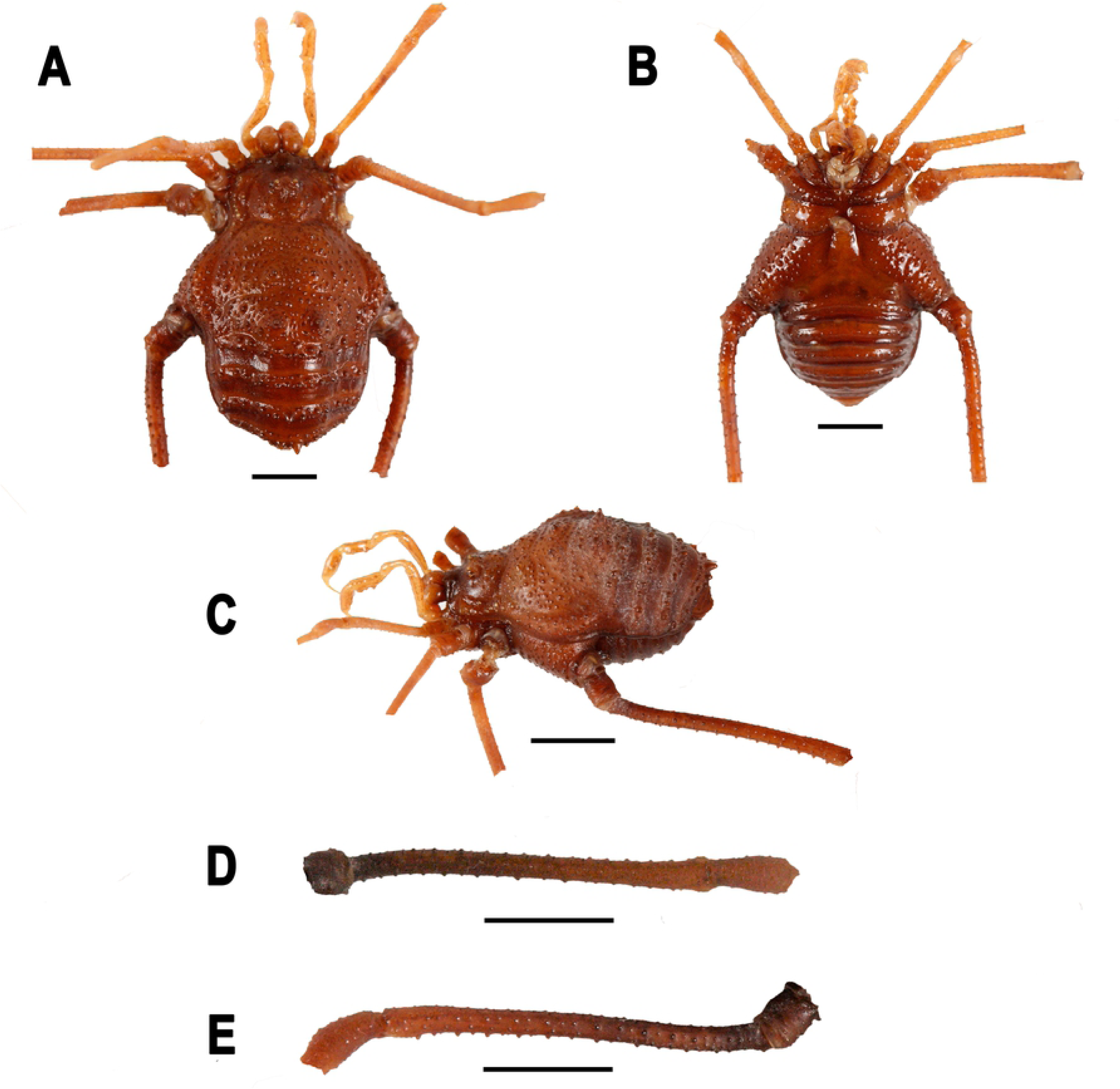
*Neosadocus robustus* (*Ilhania robusta* female paralectotype, MNRJ 42289). A. Dorsal view. B. Ventral view. C. Left lateral view. D–E. Right trochanter–patella IV (D: retrolateral view; E: prolateral view). Scale bars: 3 mm.

**Fig 8.**
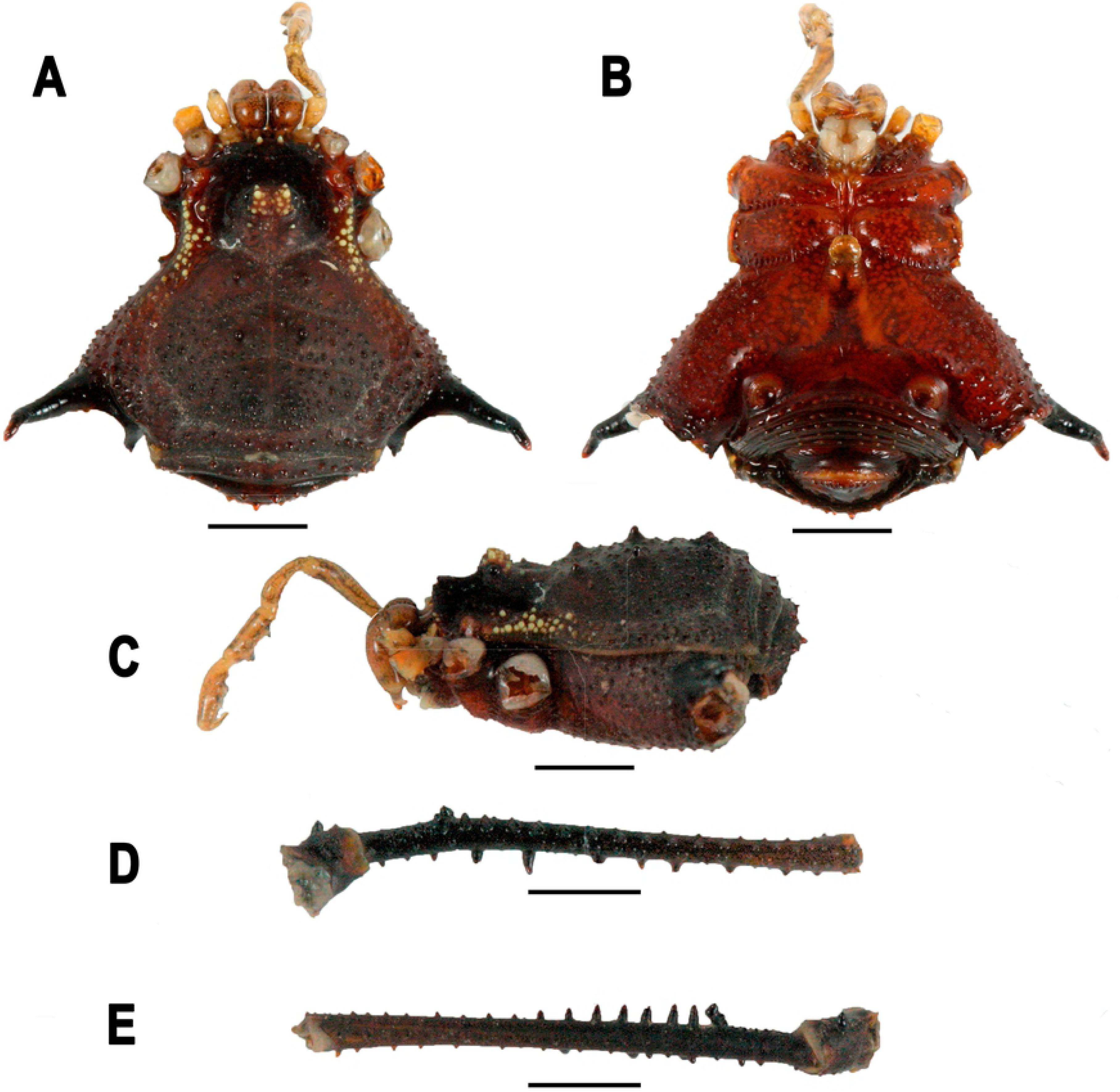
*Neosadocus maximus* (male voucher specimen, MZSP 76344). A. Dorsal view. B. Ventral view. C. Left lateral view. D–E. Right trochanter–femur IV (D: retrolateral view; E: prolateral view). Scale bars: 3 mm.

**Fig 9.**
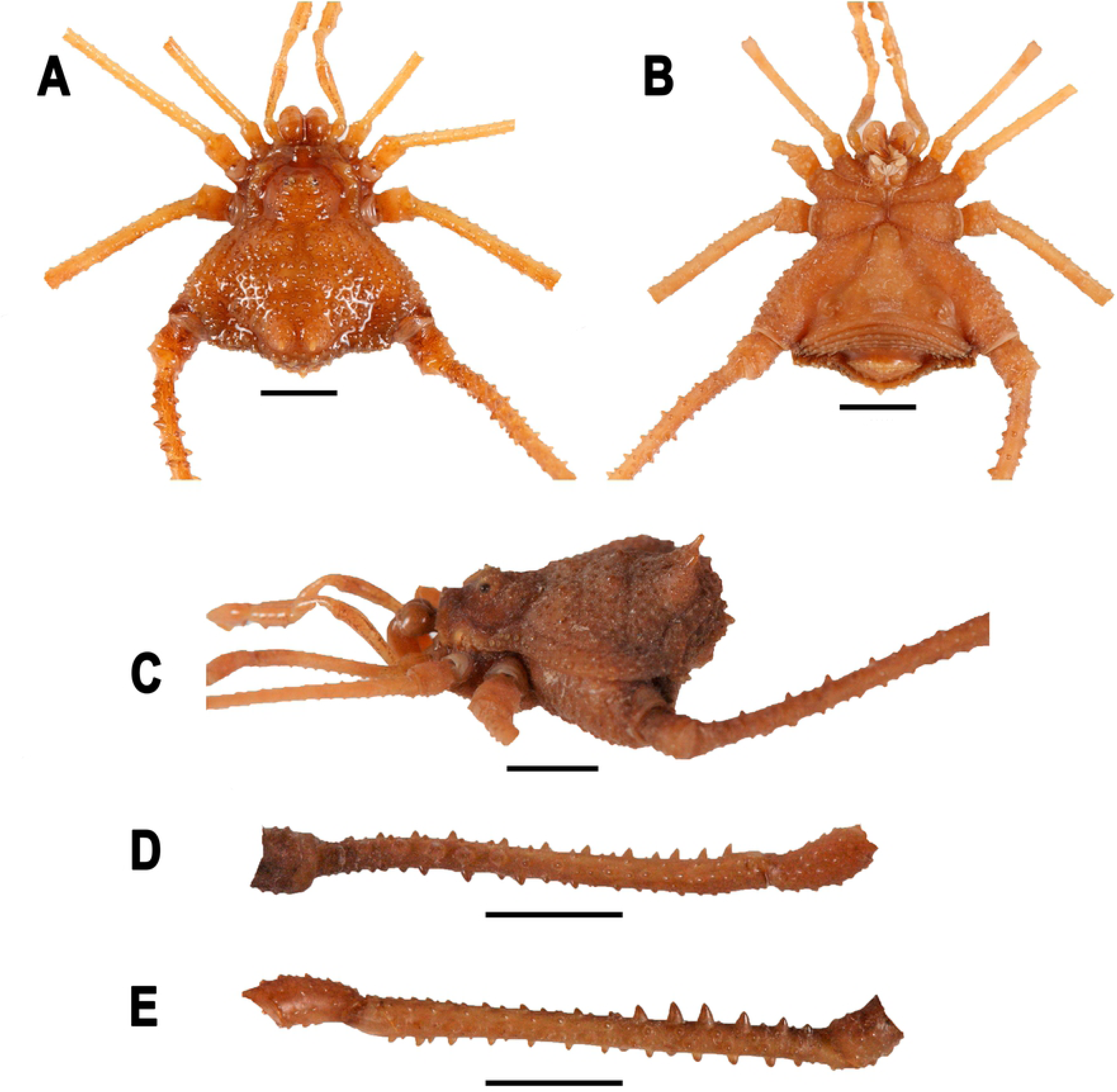
*Neosadocus maximus* (*Bunoweyhia minor* female syntype, MNRJ 41806). A. Dorsal view. B. Ventral view. C. Left lateral view. D–E. Right trochanter–patella IV (D: retrodorsal view; E: retrolateral view). Scale bars: 3 mm.

**Fig 10.**
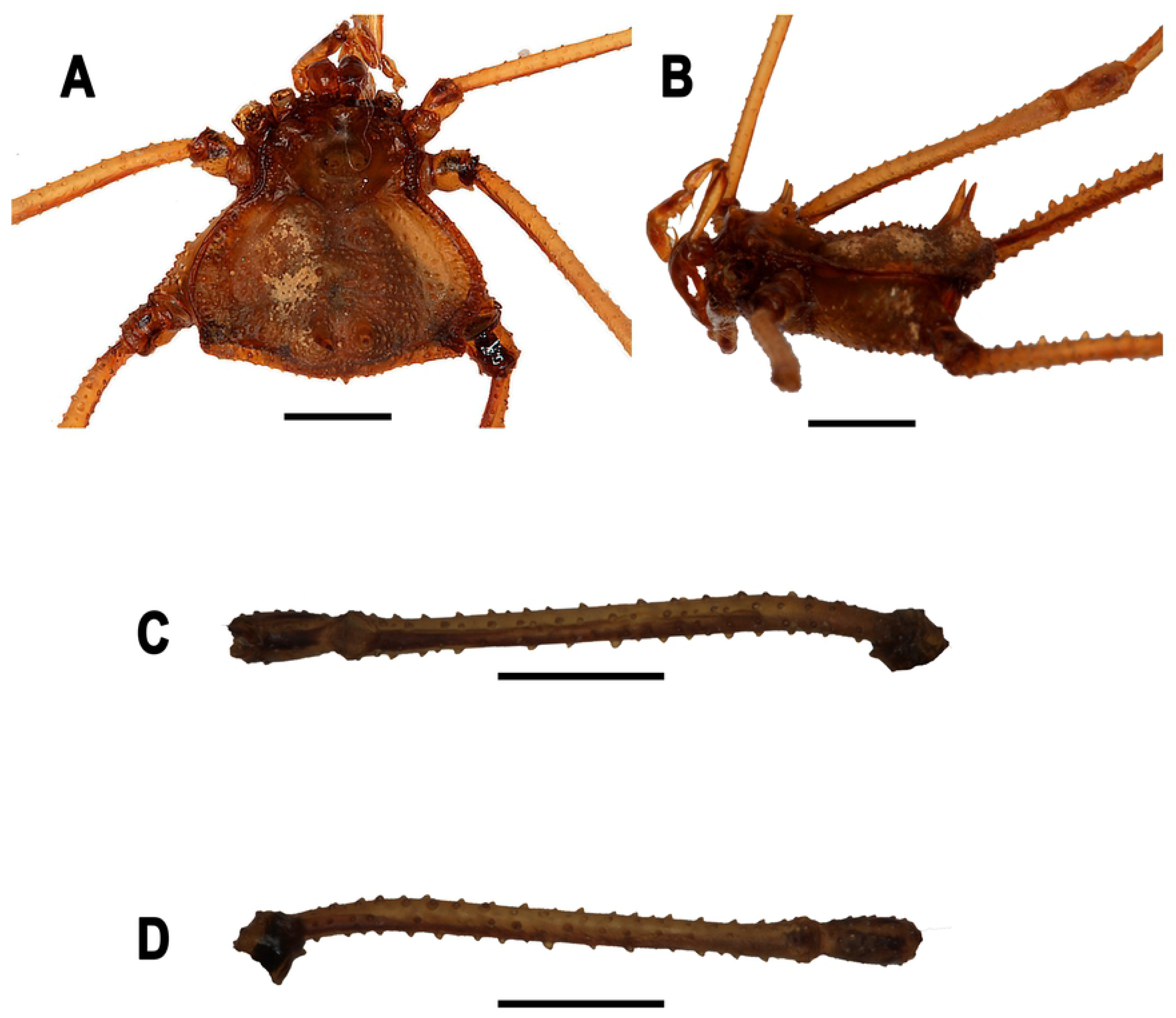
*Neosadocus misandrus* (*Metagonyleptes misandrus* female holotype, IBSP 12). A. Dorsal view. B. Left lateral view. C–D. Right trochanter–patella IV (C: retroventral view; D: dorsal view). Scale bars: 3 mm.

**Fig 11.**
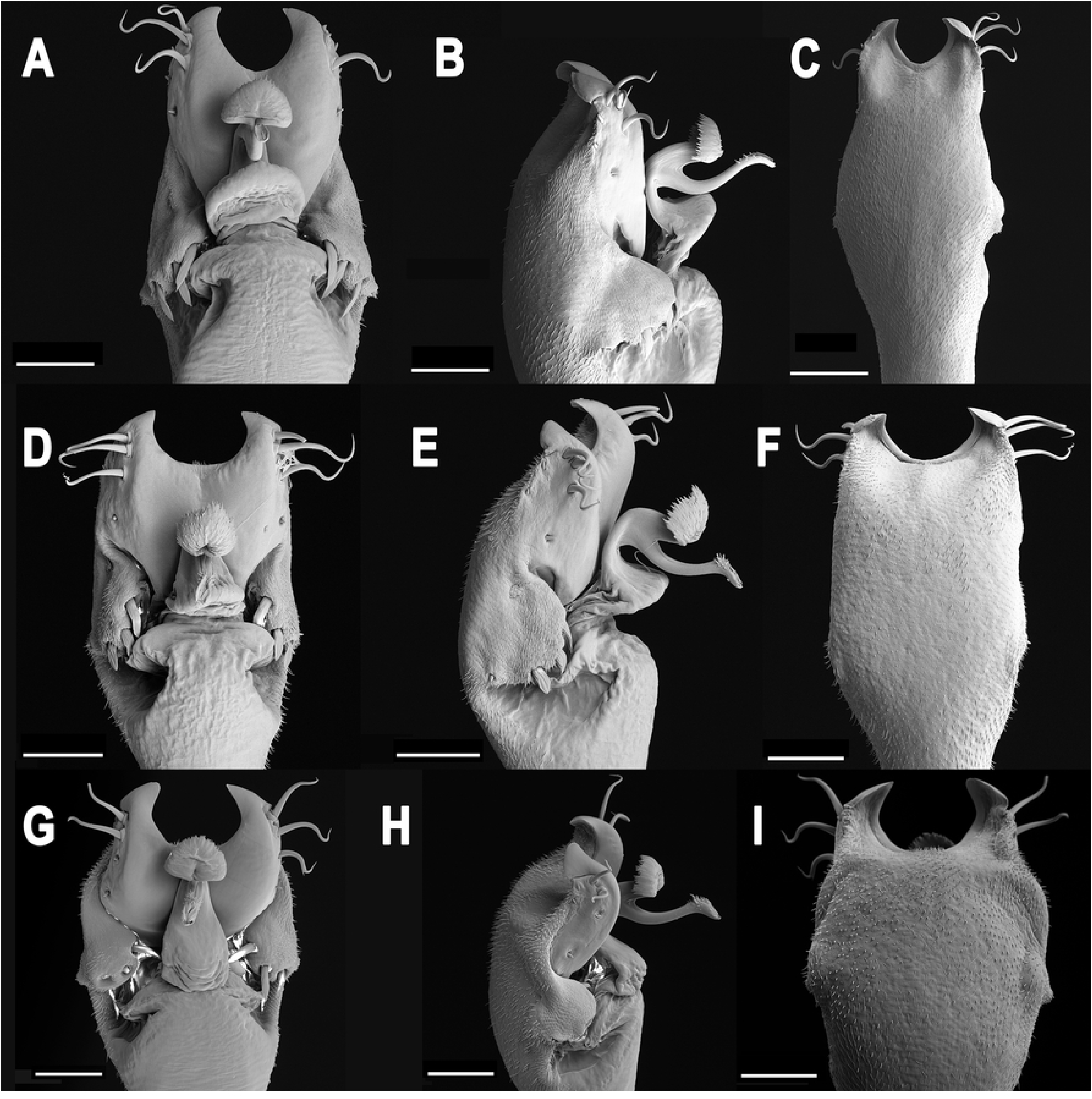
Male genitalia of *Neosadocus* species. A, D, G: dorsal view; B, E, H: left lateral view; C, F, I: ventral view. A–C. *N. bufo* (voucher specimen MZSP 76295). D–F. *robustus* (voucher specimen MZSP 76362). G–I. *N. maximus* (voucher specimen MZSP 76336). Scale bars: 100 µm.

*Sadocus* [part]: Mello-Leitão, 1923: 151 [50]; Kury, 2003: 134 [2].

*Neosadocus* Mello-Leitão, 1926: 378 [13]; Kury, 2003: 134 (complete citation listing until 2003) [2]; Machado et al., 2004: 1, 3, 5, 6, 7, 9, figs 2–2b, 11, 14, 18, 19 [51]; Willemart et al., 2007: 40, 41, fig 1, 42, fig 3, 43, fig 4, 44, figs 5 and 7, 46, 47, figs 11–12, 48, figs 13–17, 50, 51, figs 25–27, 52 [52]; Coronato-Ribeiro et al., 2013: 505 [53]; Pinto-da-Rocha et al., 2014: 527 [42]; Benedetti & Pinto-da-Rocha, 2019: 487 [54].

**Fig 12.**
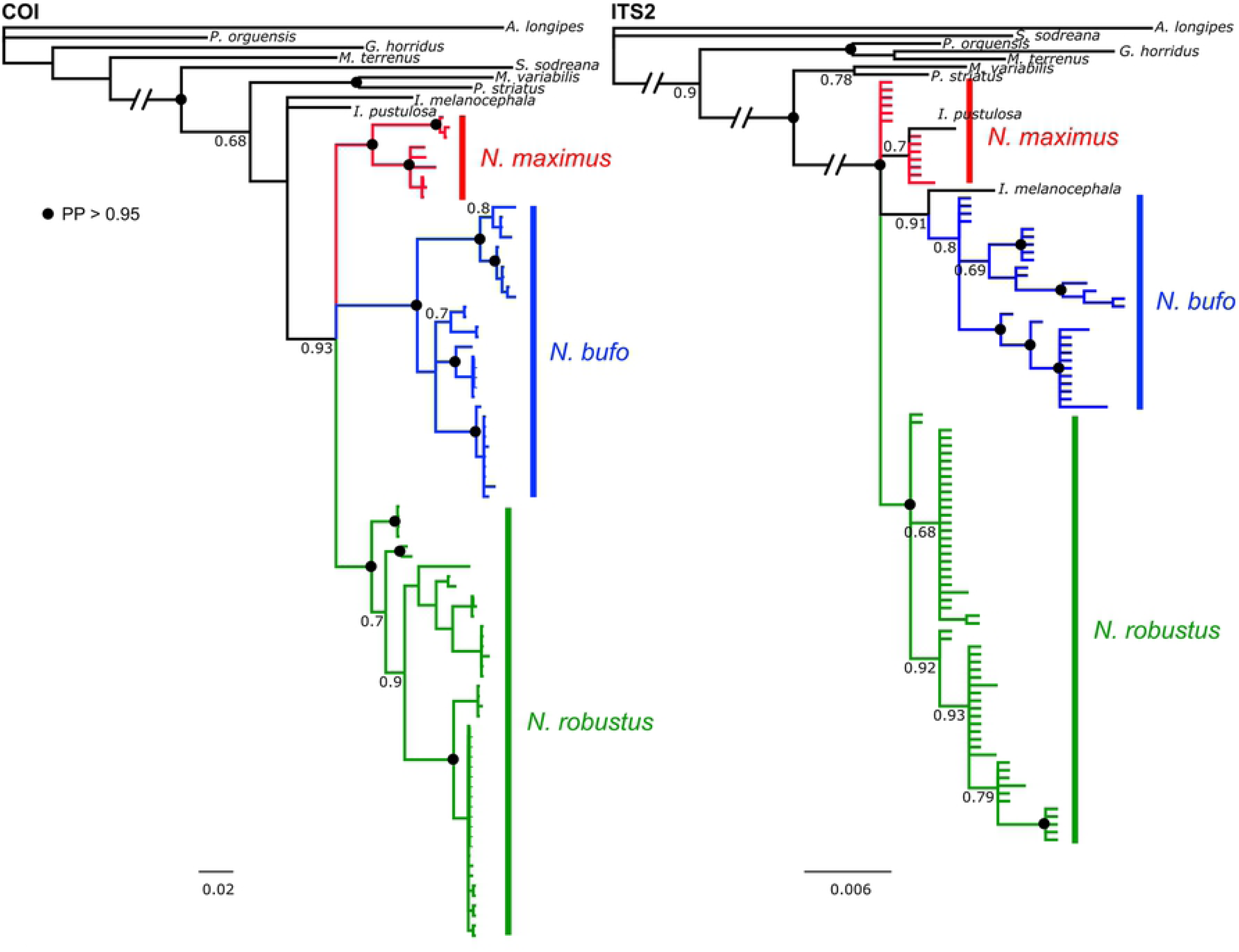
Topologies recovered under Bayesian inference analyses with MrBayes. On the left, topology inferred for COI sequences; on the right, for ITS2 sequences. In the nodes, the posterior probability values; black circles represent posterior probability *>* 0.95.

**Fig 13.**
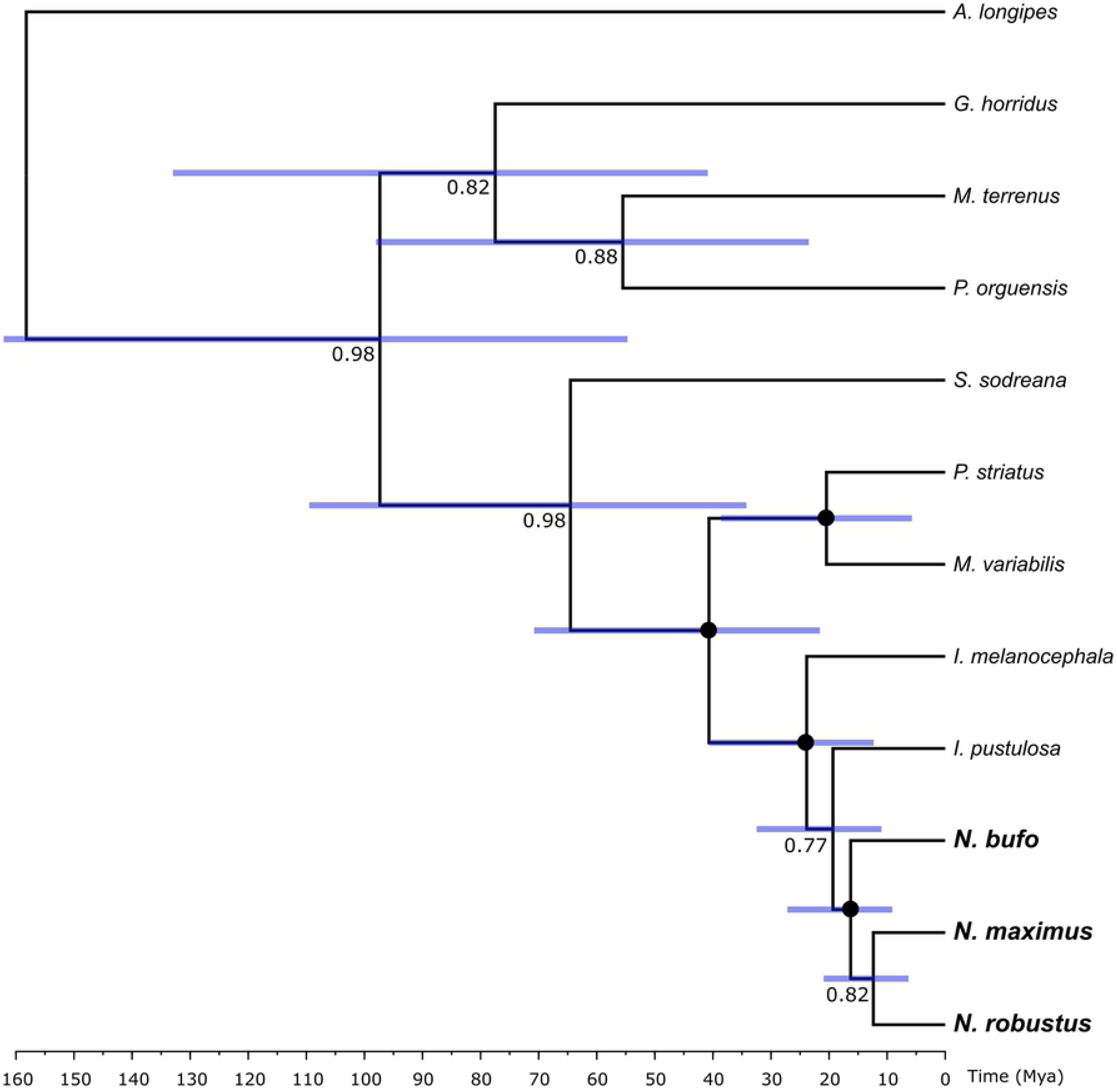
Multilocus calibrated tree obtained with *BEAST. The time scale is represented in millions of years. In the nodes, the posterior probability values; black circles represent posterior probability = 1. Blue bars show the 95% highest posterior density (HPD) intervals.

**Type species:** *Sadocus bufo* Mello-Leitão, 1923, by original designation.

*Neomitobatoides* [part]: Giltay, 1928: 84 [55]; Kury, 2003: 134 [2].

*Mitoperna* Roewer, 1931: 115 [56]; Kury, 2003: 134 (complete citation listing until 2003) [2]. Type species: *Neomitobates maximus* Giltay, 1928, by monotypy. Synonymy established by Kury, 1995 [15].

*Bunoweyhia* Mello-Leitão, 1935a: 18 [57]; Kury, 2003: 134 (complete citation listing until 2003) [2]. Type species: *Bunoweyhia variabilis* Mello-Leitão, 1935, by Kury, 2003. Synonymy established by Soares, 1944a [14].

*Ilhania* Mello-Leitão, 1936: 14 [58]; Kury, 2003: 134 [2]. Type species: *Ilhania robusta* Mello-Leitão, 1936, by original designation. Synonymy established by Soares, 1944a [14].

*Polybunos* Piza, 1943: 44 [59]; Kury, 2003: 134 [2]. Type species: *Polybunos tuberculatus* Piza, 1943, by original designation. Synonymy established by Soares, 1944a [14].

**Included species**. *Neosadocus bufo* (Mello-Leitão, 1923); *Neosadocus robustus* (Mello-Leitão, 1936); *Neosadocus maximus* (Giltay, 1928); *Neosadocus misandrus* (Mello-Leitão, 1934).

**Diagnosis.** *Neosadocus* is distinguished from other Gonyleptinae genera by the following combination of characters: <u>a. males</u>: (i) outline of dorsal scutum *γ*T-type (except in some *N. robustus* specimens with outline *γ*-type); (ii) strongly granular dorsal scutum with two blunt (*N. bufo* and *N. maximus*) or conical (*N. robustus*) paramedian tubercles on area II; (iii) two rounded (*N. bufo* and *N. maximus*) or conical (*N. robustus*) paremedian elevations on area III, both densely tuberculate; (iv) femur IV prolaterally curved (except in some *N. robustus* specimens with femur IV straight); (v) basal cluster of tubercles on femur IV; and (vi) dorso-basal apophysis of femur IV with apex curved anteriorly, followed posteriorly by a row of long tubercles. <u>b. females</u>: (i) outline of dorsal scutum *γ*-type; (ii) strongly granular dorsal scutum with two conical paramedian elevations on area III; (iii) free tergites II and III each with one central spine (with the exception of the female holotype of *N. misandrus*, which lacks a central spine on free tergites II and III); and (iv) femur IV straight, with blunt (*N. bufo*, *N. robustus* and *N. misandrus*) or conical (*N. maximus*) tubercles curved posteriorly. In both sexes, the ocularium has two conspicuous paramedian tubercles (with the exception of the female holotype of *N. misandrus*, instead with two long paramedian spines on the ocularium), and the paramedian tubercles on areas I and II are surrounded each by a circle of smaller tubercles.

### Key to the males of *Neosadocus*

**Note:** There are two types of outlines of the dorsal scutum in the males of *Neosadocus robustus* : *γ*T-type (major males) and *γ*-type (minor males). This is an important diagnostic character for the males of the species and the reason why the species is listed twice on the identification key.

1. Outline of dorsal scutum *γ*-type; proapical apophysis of coxa IV reduced in length; femur IV straight…………………………………………………………… *Neosadocus robustus*
– Outline of dorsal scutum *γ*T-type (Figs 3, 4A, 6A, 8A); proapical apophysis of coxa IV long; femur IV prolaterally curved (Figs 4D–E, 6D–E, 8D–E)…………………………………2
2. Anterior margin of carapace with three tubercles on each side (Figs 3C, 8A); femur IV with a dorso-basal apophysis of same size as the longest prodorsal tubercle near it (Figs 8D–E)……………………………………………………………… *Neosadocus maximus*
– Anterior margin of carapace with two tubercles on each side (Figs 3A–B, 4A, 6A); femur IV with a dorso-basal apophysis longer than the longest prodorsal tubercle near it (Figs 4D–E, 6D–E)…………………………………3
3. Area II with two blunt paramedian tubercles (Figs 3A, 4A and C); area III with two rounded elevations densely covered by tubercles (Figs 3, 4A and C); dorso-basal apophysis of femur IV with an apical cluster of tubercles (Figs 4D–E)……………………………………………………………………….. *Neosadocus bufo*
– Area II with two conical paramedian tubercles (Figs 3B, 6A and C); area III with two conical elevations covered by scattered tubercles (Figs 3B, 6A and C); dorso-basal apophysis of femur IV with one small basal tubercle emerging posteriorly (Figs 6D–E)………………………………………………….. *Neosadocus robustus*

### Key to the females of *Neosadocus*

1. Ocularium with long paramedian conical spines (Figs 10A–B)……………………………………………………………… *Neosadocus misandrus*
– Ocularium with two conspicuous paramedian tubercles (Figs 5A–B, 7A–B, 9A–B)…………………………………………………………………..2
2. Femur IV with rounded tubercles of similar size (Figs 7D–E). *Neosadocus robustus*
– Femur IV with dorsal row of tubercles decreasing in size distally (Figs 5D–E, 9D–E)…………………………………………………………………..3
3. Anterior margin of carapace with four tubercles on each side (Fig 5A); femur IV with blunt tubercles (Figs 5D–E)…………………………………………… *Neosadocus bufo*
– Anterior margin of carapace with three tubercles on each side (Fig 9A); femur IV with conical tubercles (Figs 9D–E)………………………………….. *Neosadocus maximus*

### *Neosadocus bufo* (Mello-Leitão, 1923)

Figs 1A–B, 3A, 4, 5, 11A–C

*Sadocus bufo* Mello-Leitão, 1923: 151, fig 23 [50]; Kury, 2003: 134 [2]. **Types** MZSP, one male holotype; MLPC, one female paratype.

*Neosadocus bufo*: Mello-Leitão, 1926: 378 [13]; Kury, 2003: 134 (complete citation listing until 2003) [2].

*Polybunos tuberculatus* Piza, 1943: 45, fig 4 [59]; Kury, 2003: 134 (complete citation listing until 2003) [2]. Type MZSP 66, one female holotype. Synonymy established by Soares, 1944a [14].

*Bunoweyhia lata* Mello-Leitão, 1935b: 390, fig 18 [60]; Kury, 2003: 134 (complete citation listing until 2003) [2]. Type MNRJ 42364, one male holotype. Synonymy established by Soares & Soares, 1984 [61].

*Neosadocus latus*: Soares, 1945: 361 [62]; Kury, 2003: 134 (complete citation listing until 2003) [2].

*Bunoweyhia variabilis* Mello-Leitão, 1935a: 18, figs 10–10a [57]; Kury, 2003: 134 (complete citation listing until 2003) [2]. Type MNRJ 41803, one male and one female syntypes. **SYNONYMY RESTORED**, as performed by Soares & Soares, 1984 [61].

*Neosadocus variabilis*: Soares, 1944b: 282 [63]; Kury, 2003: 135 (complete citation listing until 2003) [2].

**Type material.** *Sadocus bufo* Mello-Leitão, 1923 (MZSP, one male holotype; MLPC, one female paratype). Locality: Brazil, Rio de Janeiro, Petrópolis (NOT EXAMINED). *Polybunos tuberculatus* Piza, 1943 (MZSP 66, one female holotype). Locality: Brazil, São Paulo, Itapecerica: Bat^ea (EXAMINED). *Bunoweyhia lata* Mello-Leitão, 1935 (MNRJ 42364, one male holotype). Locality: Brazil, São Paulo, Lussanvira (EXAMINED). *Bunoweyhia variabilis* Mello-Leitão, 1935 (MNRJ 41803, one male and one female syntypes). Locality: Brazil, São Paulo, Ribeira do Iguape (EXAMINED).

**Other material examined.** BRAZIL. <u>São Paulo</u>: Cotia (Reserva Florestal Morro Grande), 1F (MZSP 25969); ibidem, 1F (MZSP 30331); ibidem, 1F (MZSP 30332); ibidem, 1M (MZSP 30349); ibidem, 1M (MZSP 30345); ibidem, 4M 2F (MZSP 30346); ibidem, 1M (MZSP 30351); ibidem, 1M 1F (MZSP 30352); ibidem, 2F (MZSP 25970); ibidem, 1M (MZSP 25971); ibidem (Capelinha), 1M (MZSP 25964); ibidem, 1M (MZSP 25965); ibidem (Grilos), 1M (MZSP 25966); ibidem, 4M 3F (MZSP 27913); ibidem, 1F (MZSP 25967); ibidem, 1M 1F (MZSP 27914); ibidem, 1M (MZSP 27915); ibidem, 3M 1F (MZSP 27916); ibidem, 6M 5F (MZSP 27917); ibidem, 6M 5F (MZSP 27918); ibidem, 4M 1F (MZSP 27919); ibidem (Quilombo), 2F (MZSP 25955); ibidem, 2F (MZSP 25956); ibidem, 2M 2F (MZSP 25957); ibidem, 1F (MZSP 25960); ibidem, 1M (MZSP 25962); ibidem, 1M 1F (MZSP 25963); ibidem, 1M (MZSP 76332); ibidem, 1M (MZSP 76333); ibidem, 1M (MZSP 76334); ibidem, 2M 1F (MZSP 30348); ibidem, 1M (MZSP 27896); ibidem, 1M (MZSP 27897); ibidem, 1M (MZSP 27899); ibidem, 1M (MZSP 27900); ibidem, 7M 1F (MZSP 27901); ibidem, 1M 2F (MZSP 27902); ibidem, 1M 1F (MZSP 27903); ibidem, 1M 1F (MZSP 27905); ibidem, 2M (MZSP 28904); ibidem (Torres), 2M 3F (MZSP 27920); Guapiara (Fazenda Intervales), 4M 1F (MZSP 14096); ibidem (Ouro Grosso), 1M (MZSP 42066); ibidem (Trilha Lago Negro, Fazenda Intervales), 1M 2F (MZSP 14409); Iporanga, 1M 2F (MZSP 11357); Picinguaba, 1F (MZSP 19136); Pilar do Sul (Fazenda do Linho), 1F (MZSP 21582); Sete Barras (Saibadela, Fazenda Intervales), 2F (MZSP 16745); Cajati (Barra do Azeite, Cachoeira do Azeite), 1F (MZSP 76324); ibidem, 1F (MZSP 76325); ibidem, 1M (MZSP 76326); Canańeia (Ilha do Cardoso), 4F (MZSP 29318); Capão Bonito (Parque Estadual Intervales), 7M 4F (MZSP 30091); Iguape (km 236 of the road, near Sete Barras municipality), 1M (MZSP 42070); Iguape (Rodovia Prefeito Casemiro Teixeira), 1F (MZSP 76302); ibidem, 1F (MZSP 76303); ibidem, 1M (MZSP 76304); ibidem, 1M (MZSP 76305); ibidem, 1M (MZSP 76306); ibidem, 1M (MZSP 76307); ibidem, 1M (MZSP 76308); ibidem, 1M (MZSP 76309); ibidem, 1F (MZSP 76310); Iporanga (Lajeado), 3M 1F (MZSP 832); ibidem, 1F (MZSP 843); Iporanga (Parque Estadual Tuŕıstico do Alto Ribeira, Santana) 4M 4F (MZSP 26820); Iporanga (Pousada Rupestre), 1M (MZSP 76327); ibidem, 1M (MZSP 76328); ibidem, 1M (MZSP 76329); Itapecerica da Serra (Bat^ea), 1M (MZSP 67); ibidem, 1M 1F (MZSP 879); Juquiá (Hotel Fazenda Dois Palmitos), 1F (MZSP 76330); ibidem, 1M (MZSP 76331); Miracatu (Cachoeira do Faú), 1M (MZSP 76301); ibidem (Rodovia Ŕegis Bittencourt, km 347), 1M (MZSP 76311); ibidem, 1F (MZSP 76312); ibidem, 1M (MZSP 76313); ibidem, 1M (MZSP 76314); ibidem, 1M (MZSP 76315); ibidem, 1M (MZSP 76316); ibidem, 1F (MZSP 76317); ibidem, 1M (MZSP 76318); ibidem, 1M (MZSP 76319); ibidem, 1M (MZSP 76320); ibidem, 1M (MZSP 76321); ibidem, 1F (MZSP 76322); ibidem, 1F (MZSP 76323); Perúıbe, 1M (MZSP 29946); Poço Grande, 5F (MZSP 1633); Ribeirão Grande (Parque Estadual Intervales), 1M (MZSP 76292); ibidem, 1M (MZSP 76293); ibidem, 1M (MZSP 76294); ibidem, 1M (MZSP 76295); ibidem, 1M (MZSP 76296); ibidem, 1M (MZSP 76297); ibidem, 1M (MZSP 76298); ibidem, 1M (MZSP 76299); ibidem, 1M (MZSP 76300); ibidem, 1M 2F (MZSP 30212); ibidem, 1M (MZSP 29898); ibidem, 3M (MZSP 30096); São Miguel Arcanjo (Parque da Onça Parda), 2M 2F (MZSP 46706); ibidem (Parque Estadual Carlos Botelho), 2M 1F (MZSP 45551); São Vicente (Japúı, trail to Praia de Itaquitanduva), 6M 6F (MZSP 29938).

**Distribution (Fig 2).** BRAZIL: Southern coast of the state of São Paulo. There are two questionable records of *N. bufo*: one record from the town of Petrópolis (Southeastern region of Rio de Janeiro state), the locality of the type-species of the genus, *Sadocus bufo*. We did not not analyze this specimen because it is lost; the other record belongs to Lussanvira (Northwestern region of the state of São Paulo), the type locality of *Polybunos tuberculatus*. Despite the fact that a great number of Opiliones have been collected in both localities, *Neosadocus* specimens have not been found among them.

**Diagnosis.** *Neosadocus bufo* males differ from the other males of *Neosadocus* in having dorsal scutum wider than long. Also, *N. bufo* males possess an apical cluster of tubercles on the anterior surface of the dorso-basal apophysis of femur IV. Otherwise, *N. robustus* and *N. maximus* males exhibit small sparse tubercles, not forming a cluster. *Neosadocus bufo* females differ from the other *Neosadocus* females in having two lateral clusters of four small tubercles on both sides of the anterior margin of the carapace. Additionally, the paramedian spines on area III of *N. bufo* females measure over 2.0 mm, longer than in the females of the other species.

Redescription of the male (*Bunoweyhia variabilis* male syntype, MNRJ 41803).

*Measurements* : **DSL**: 9.1 mm. **DSW**: 10.3 mm. **LDA**: 1.5 mm.

*Dorsum* (Figs 3A, 4A and C): **Outline of dorsal scutum** *γ*T-type. **Dorsal scutum** wider than long. **Anterior margin of carapace** with two small tubercles on each side. **Ocularium** tuberculate, with two conspicuous paramedian tubercles. **Carapace area behind the ocularium** tuberculate. **Lateral margin of prosoma** with tubercles concentrated on the posterior half region. **Areas I–II** tuberculate, each with two prominent blunt paramedian tubercles surrounded by a circle of smaller tubercles. **Area III** tuberculate, with two rounded paramedian elevations densely tuberculate, each with one central tubercle. **Lateral margin of dorsal scutum** tuberculate, with an organized external row of tubercles. **Free tergites I–III** unarmed and with tubercles of different sizes.

*Venter* (Fig 4B): **Coxa I** with a central longitudinal row of four prominent conical tubercles and small sparse tubercles; with a cluster of three prominent tubercles and one isolated prominent tubercle on distal margin. **Coxa II** with three central prominent conical tubercles and small sparse tubercles; with a cluster of three prominent tubercles and a cluster of two prominent tubercles on distal margin. **Coxa III** with small sparse tubercles; with an anterior row of eight prominent flattened tubercles and a posterior row of 12 prominent flattened tubercles, the latter almost fused with coxa IV. **Coxa IV** tuberculate. **Stigmatic area** tuberculate, with a posterior row of small tubercles. **Genital operculum** with small sparse tubercles. **Sternites I–IV** each with a row of small tubercles. **Sternite V** tuberculate. **Anal operculum** with sparse tubercles of different sizes.

*Chelicera*: **Segment I** with a cluster of three small projections and one small prolateral projection on distal margin. **Bulla** with small sparse ventral tubercles and nine spiniform ventral tubercles on distal margin. **Segment II** with 10 teeth. **Segment III** with six teeth, higher than those of segment II.

*Pedipalp*: **Coxa** with a cluster of seven ventro-apical tubercles and one retrodorsal apical tubercle. **Trochanter** with small sparse dorsal tubercles and two ventro-apical tubercles. **Femur** elongated, with small sparse dorsal tubercles and a ventral row of six small sparse tubercles. **Patella** with small setae both prolaterally and retrolaterally on distal margin; with one small prolateral tubercle and one small retrolateral tubercle on proximal margin; with two small ventral tubercles on proximal margin. **Tibia** with small sparse setae; tibial setation: prolateral IiIi and retrolateral IIi. **Tarsus** with small sparse setae; tarsal setation: prolateral IiIiiii and retrolateral IIiii.

*Legs* (Figs 4D–E show leg IV): **Coxae I–III** each with one prodorsal and one retrodorsal tubercle. **Coxa III** with a retrolateral row of tubercles. **Coxa IV** tuberculate, with a long proapical apophysis. **Proapical apophysis of coxa IV** with small dorsal tubercles and sparse ventral setae; with one retroventral proximal tubercle and three distal inflated ventral projections. **Trochanters I–III** tuberculate, each with one prodorsal and one retroventral tubercle on distal margin. **Trochanters I–II** longer than wide, each with one prodorsal and one retroventral tubercle on proximal margin. **Trochanter III** as wide as long. **Trochanter IV** wider than long and sparsely tuberculate; prolaterally with one conspicuous blunt central tubercle and one proximal cluster of seven small setae; retroventrally with one conspicuous apical tubercle. **Femora I–IV** with approximately six longitudinal rows (prodorsal, retrodorsal, retrolateral, retroventral, proventral and prolateral). **Femora I–II** straight, unarmed, with small sparse tubercles. **Femur II** with tubercles near the distal margin subtly longer. **Femur III** with sparse conical tubercles of different sizes and two curved ventral tubercles on distal margin. **Femur IV** prolaterally curved; proximally with small sparse retrolateral tubercles, one dorsal cluster of 32 small tubercles, one ventral cluster of three tubercles and one isolated ventral tubercle; distally with a dorsal cluster of seven small tubercles and two ventral spines, all curved posteriorly; prodorsal and retrodorsal rows of tubercles each decreasing in size distally (the three basal tubercles of prodorsal row longer, the basalmost tubercle almost fused with a dorso-basal apophysis); retrolateral row of tubercles decreasing in size distally (the four basal tubercles large); retroventral and prolateral rows with small tubercles; proventral row with tubercles of different sizes, almost interspersed; the distalmost tubercles of each row curved posteriorly. **Dorso-basal apophysis of femur IV** inclined distally, with the apex curved anteriorly; 1.5x longer than the longest prodorsal tubercle near it and with small tubercles on the anterior face; with an apical cluster of three tubercles. **Patellae I–IV** unarmed. **Patellae I–II** with small sparse tubercles. **Patellae III–IV** with sparse tubercles curved posteriorly. **Tibiae I–IV** unarmed. **Tibiae I–II** smooth. **Tibiae III–IV** with tubercles curved posteriorly. **Tibia III** shorter than the other tibiae and with two ventral rows of tubercles increasing in size distally. **Tibia IV** with one ventral row of tubercles increasing in size distally. **Tarsal formula**: 6(3), 12(3), 7, 8.

*Coloration in ethanol* (voucher specimen MZSP 76313): **Ocularium, carapace area behind the ocularium, areas I–III, lateral margin of dorsal scutum, trochanters I–III, venter of coxae I–III, coxa IV, tubercles of area I and coxa IV, genital operculum, stigmatic area, and ventral tubercles** strong orange yellow (centroid 68). **Anterior margin of carapace, carapace area between ocularium and lateral margin of prosoma, dorsum of coxae I–III, proapical apophysis of coxa IV, dorsum of trochanter IV, tubercles of areas II–III, sternites I–V, and anal operculum** brownish black (centroid 65). **Chelicerae, pedipalps, femur I, patellae I–II, tibiae I–II, metatarsi I–IV, and tarsi I–IV** brilliant greenish yellow (centroid 98). **Posterior margin of dorsal scutum, free tergites I–III, femora II–IV, patellae III–IV, tibiae III–IV, and venter of trochanter IV** brownish orange (centroid 54). **Tubercles of frontal hump, of ocularium and of lateral margin of prosoma** pale yellow (centroid 89). This coloration pattern is also observed in living specimens.

*Genitalia* (voucher specimen MZSP 76295; Figs 11A–C): **Glans** flattened and wrinkled. **Stylus** elongated and curved, with the apex dorsally projected and with subapical ventral trichomes. **Ventral process** half the stylus length and ventrally concave, with various trichomes on the margin and on dorsal face. **Ventral plate** subrectangular and with a U-shaped apex; with three pairs of macrosetae A, one pair of macrosetae B, three pairs of macrosetae C, one pair of macrosetae D and two pairs of macrosetae E.

*Variation in males* (n = 28): **Femur IV** with 1–3 long tubercles near the dorso-basal apophysis. **Dorsal scutum coloration** brownish black (centroid 65) with patches between strong orange (centroid 50) and strong orange yellow (centroid 68). **Leg coloration** from brownish black (centroid 65) to deep orange (centroid 51).

Redescription of the female (*Bunoweyhia variabilis* female syntype, MNRJ 41803).

*Measurements* : **DSL**: 9.5 mm. **DSW**: 8.0 mm. **LPA**: 2.5 mm.

*Dorsum* (Figs 5A and C): **Outline of dorsal scutum** *γ*-type. **Dorsal scutum** longer than wide. **Frontal hump** with two paramedian tubercles. **Anterior margin of carapace** with four small tubercles on each side. **Area III** with two long paramedian conical spines encompassed by various tubercles. **Free tergites II–III** each with one central spine.

*Venter* (Fig 5B): **Coxa I** with a proximal cluster of three prominent tubercles and a distal cluster of two prominent tubercles. **Coxa II and stigmatic area** tuberculate. *Pedipalp*: **Femur** smooth. **Tibial setation**: prolateral Ii(Ii) and retrolateral IIi.

**Tarsal setation**: prolateral IiIiiiii and retrolateral IIiii.

*Legs* (Figs 5D–E show leg IV): **Coxa III** smooth. **Coxa IV** shorter than in males; proapical apophysis absent. **Trochanter IV** without a prolateral apophysis; retroventral tubercle and prolateral proximal cluster of small setae both absent. **Femora II–III** tuberculate. **Femur IV** straight, with blunt tubercles decreasing in size distally, all curved posteriorly; dorso-basal apophysis absent. **Tarsal formula**: 3(3), 7(3), 7, 8.

Variation in females not detected.

### *Neosadocus robustus* (Mello-Leitão, 1936)

Figs 1C–D, 3B, 6, 7, 11D–F, S1 Fig

*Ilhania robusta* Mello-Leitão, 1936: 14, fig 11 [58]; Kury, 2003: 135 [2]. **Type** MNRJ 42289, one male lectotype and one female paralectotype, here designated.

*Neosadocus robustus* : Soares, 1945: 217 [64]; Kury, 2003: 135 [2].

*Neosadocus bufo* [part]: Soares, 1944a: 244 [14]; Kury, 2003: 135 [2].

**Type material.** *Ilhania robusta* Mello-Leitão, 1936 (MNRJ 42289, one male lectotype and one female paralectotype, here designated). Locality: Brazil, Parańa, Antonina (EXAMINED).

**Other material examined.** BRAZIL. <u>Parańa</u>: Antonina (Pousada Curupira), 1F (MZSP 76393); ibidem, 1M (MZSP 76394); ibidem, 1M (MZSP 76395); ibidem, 1F (MZSP 76396); ibidem, 1M (MZSP 76397); ibidem, 1M (MZSP 76398); ibidem (Praia Grande), 1F (MZSP 36337); ibidem, 1M (MZSP 36348); ibidem (Trilha do Rio Xaxim), 1M (MZSP 76390); ibidem, 1M (MZSP 76391); ibidem, 1F (MZSP 76392); Curitiba, 4F (MZSP 17123); ibidem, 1M (MZSP 30200); ibidem, 2F (MZSP 36424); ibidem (Merĉes), 1M 1F (MZSP 36371); ibidem (Pilarzinho), 1F (MZSP 36336); ibidem (Parque Barigui), 2M 1F (MZSP 770); ibidem, 1M 2F (MZSP 918); Fazenda Rio Grande, 1M (MZSP 76365); Guaraqueçaba (Rio Poruquara), 1M (MZSP 15991); ibidem, 1M 2F (MZSP 17106); ibidem (Parque Nacional do Superagui), 2M (MZSP 30097); ibidem (Reserva Natural Rio Cachoeira), 1F (MZSP 76403); ibidem (Tagagaba, 150m high), 1F (MZSP 76379); ibidem, 1F (MZSP 76380); ibidem, 1M (MZSP 76381); ibidem, 1M (MZSP 76382); 1M (MZSP 76383); ibidem (Trilha do Ferro), 1M (MZSP 76399); ibidem, 1M (MZSP 76400); 1M (MZSP 76401); ibidem, 1F (MZSP 76402); Guaratuba (Pedra Branca de Araraquara, Rio Bonito), 1F (MZSP 76356); ibidem (Rio Bonito, BR101 km 680), 2M 1F (MZSP 29398); ibidem (Usina de Guaricana), 4M 2F (MZSP 18109); ibidem, 8M 5F (MZSP 18129); Morretes, 1F (MZSP 76350); ibidem, 1F (MZSP 76351); ibidem, 1F (MZSP 76352); ibidem, 1M (MZSP 76353); ibidem, 1M (MZSP 76354); ibidem (Anhaia), 1M (MZSP 36420); ibidem (Caminho do Itupava), 1F (MZSP 76357); ibidem, 1F (MZSP 76358); ibidem, 1M (MZSP 76359); ibidem, 1F (MZSP 76360); ibidem, 1F (MZSP 76361); ibidem (Marumbi), 1F (MZSP 36315); ibidem, 1F (MZSP 36331); ibidem, 1F (MZSP 36338); ibidem, 1F (MZSP 36355); ibidem, 6M (MZSP 36400); ibidem, 1M (MZSP 36408); ibidem, 3M 3F (MZSP 36412); ibidem, 1M (MZSP 36417); ibidem, 1M 1F (MZSP 36429); ibidem (Rio Taquaral), 1M 1F (MZSP 36375); ibidem (Parque Estadual do Marumbi, Caminho do Itupava), 1M (MZSP 59928); ibidem (Parque Nacional Saint Hilaire/Lange), 1F (MZSP 76355); ibidem (Porto de Cima), 1F (MZSP 76363); ibidem, 1F (MZSP 76364); Paranagúa (Parque Nacional Saint-Hilaire/Lange, Fazenda Niterói, 80m high), 1F (MZSP 76384); ibidem, 1M (MZSP 76385); ibidem, 1M (MZSP 76386); Piraquara (Banhado), 1M 1F (MZSP 898); ibidem, 1M (MZSP 908); ibidem, 1F (MZSP 36318); ibidem, 1M (MZSP 36327); ibidem, 1M (MZSP 36363); ibidem, 1M 1F (MZSP 36364); ibidem, 1M 1F (MZSP 36395); São Jośe dos Pinhais (Centro), 1M (MZSP 29403); São Jośe dos Pinhais (Usina Hidreĺetrica de Guaricana’s dam), 1M 2F (MZSP 16782); ibidem, 1M (MZSP 16783); Tunas do Paraná (near Gruta dos Jesúıtas), 2M 1F (MZSP 30223). <u>São Paulo</u>: Cotia (Reserva Florestal Morro Grande), 1F (MZSP 25972); ibidem (Reserva Florestal Morro Grande, Quilombo), 1M (MZSP 25958); ibidem, 1F (MZSP 25959); Guapiara (Bocaina, Fazenda Intervales), 3M 1F (MZSP 16781); ibidem (Cachoeira do Mirante, Fazenda Intervales), 4M 1F (MZSP 14027); Guapiara, 1M (MZSP 19282); Pilar do Sul (Fazenda Fibrasil), 3M 4F (MZSP 17150); Barra do Turvo (Parque Estadual do Rio Turvo, Núcleo Cedro), 1M (MZSP 76372); ibidem, 1F (MZSP 76373); ibidem, 1F (MZSP 76374); ibidem, 1F (MZSP 76375); ibidem, 1F (MZSP 76376); ibidem, 1F (MZSP 76377); ibidem, 1F (MZSP 76378); Cajati (Parque Estadual do Rio Turvo, Núcleo Capelinha), 1F (MZSP 76366); ibidem, 1F (MZSP 76367); ibidem, 1F (MZSP 76368); ibidem, 1M (MZSP 76369); ibidem, 1M (MZSP 76370); Cajati (Parque Estadual do Rio Turvo, near Gruta do Lamarca), 1F (MZSP 76371); Canańeia (Ilha do Cardoso), 1M 1F (MZSP 29323); ibidem, 1M 5F (MZSP 29324); ibidem, 1M 1F (MZSP 29325); ibidem, 1M (MZSP 29332); ibidem, 1M 4F (MZSP 29333); Canańeia (Serra do Itapitangui, road to Ariri, km 9), 2M 4F (MZSP 29397); ibidem (IO-USP building), 1M (MZSP 76362); Cotia (Reserva Florestal Morro Grande), 4M 3F (MZSP 27726); ibidem, 4M 1F (MZSP 30131); ibidem (Portal do Quilombo), 1M (MZSP 76413); ibidem, 1M (MZSP 76414); ibidem, 1M (MZSP 76415); ibidem, 1M (MZSP 76416); ibidem, 6M (MZSP 27732); ibidem, 1F (MZSP 27763); ibidem, 1M (MZSP 27765); ibidem, 2M 2F (MZSP 27904); ibidem, 2M 3F (MZSP 27744); ibidem, 1F (MZSP 30261); ibidem, 1F (MZSP 30265); ibidem, 1F (MZSP 30275); Cotia (Caucaia do Alto), 1F (MZSP 42081); ibidem, 1M (MZSP 42089); Ibiúna, 1M (MZSP 76387); ibidem, 1M (MZSP 76388); ibidem, 1F (MZSP 76389); Iguape (Ilha do Cardoso), 6M (MZSP 30130); Jacupiranga, 1M (MZSP 29340); ibidem (Parque Estadual Jacupiranga), 1M (MZSP 29940); Poço Grande, 4M (MZSP 1634); Ribeirão Grande (Waterfall trail at Maciel), 1M (MZSP 76404); ibidem, 1M (MZSP 76405); ibidem, 1M (MZSP 76406); ibidem, 1M (MZSP 76407); ibidem, 1M (MZSP 76408); ibidem, 1M (MZSP 76409); ibidem, 1M (MZSP 76410); ibidem, 1M (MZSP 76411); ibidem, 1M (MZSP 76412); São Miguel Arcanjo (Parque Estadual Carlos Botelho), 1M (MZSP 48262); ibidem, 1M 1F (MZSP 48263); ibidem, 4M 4F (MZSP 45593); ibidem, 1F (MZSP 45525).

**Distribution (Fig 2).** BRAZIL: From Southern coast of the state of Paraná to the Central coast of the state of São Paulo.

**Diagnosis.** *Neosadocus robustus* males differ from the other males of *Neosadocus* in having a conspicuous dorso-basal apophysis on femur IV, twice longer than the longest prodorsal tubercle near it. Also, *N. robustus* males have conspicuous conical tubercles near the dorso-basal apophysis of femur IV. Moreover, *N. robustus* is the species with the greatest morphological variation in males (S1 Fig; see Variation in males). *Neosadocus robustus* females differ from the other *Neosadocus* females by presenting two small lateral tubercles on both sides of the anterior margin of carapace. Also, *N. robustus* females have rounded tubercles of similar size on femur IV. Both males and females of *N. robustus* have anterior and posterior rows of triangular tubercles on the venter of coxa III, whereas the tubercles of males and females of both *N. bufo* and *N. maximus* are flattened.

Redescription of the male (*Ilhania robusta* male lectotype, MNRJ 42289).

*Measurements* : **DSL**: 11.1 mm. **DSW**: 11.1 mm. **LDA**: 1.9 mm.

*Dorsum* (Figs 3B, 6A and C): **Outline of dorsal scutum** *γ*T-type. **Dorsal scutum** as wide as long. **Anterior margin of carapace** with two small tubercles on each side. **Ocularium** tuberculate, with two conspicuous paramedian tubercles. **Carapace area behind the ocularium** tuberculate. **Lateral margin of prosoma** with tubercles concentrated on the posterior half region. **Area I** tuberculate, with two prominent blunt paramedian tubercles surrounded by a circle of smaller tubercles. **Area II** tuberculate, with two prominent conical paramedian tubercles surrounded by a circle of smaller tubercles. **Area III** tuberculate, with two conical paramedian elevations densely tuberculate, each with one central tubercle. **Lateral margin of dorsal scutum** tuberculate, with an organized external row of tubercles. **Free tergite I** unarmed and with tubercles of different sizes. **Free tergites II–III** each with one central spine and with tubercles of different sizes.

*Venter* (Fig 6B): **Coxa I** with a central longitudinal row of three prominent conical tubercles and one isolated prominent central conical tubercle; with small sparse tubercles and two conical tubercles on proximal margin; with a cluster of three and a cluster of two prominent tubercles on distal margin. **Coxa II** with five central prominent cone-shaped tubercles and small sparse tubercles; with a cluster of two prominent tubercles and one isolated prominent tubercle on distal margin. **Coxa III** with small sparse tubercles, with an anterior row of five prominent triangular tubercles and a posterior row of 10 prominent triangular tubercles, the latter almost fused with coxa IV. **Coxa IV** tuberculate. **Stigmatic area** sparsely tuberculate. **Genital operculum** tuberculate. **Sternites I–IV** each with a row of small tubercles.

**Sternite V and anal operculum** tuberculate.

*Chelicera* (voucher specimen MZSP 76388): **Segment I** with one small tubercle and one small prolateral projection on distal margin. **Bulla** tuberculate, with six small ventral tubercles on distal margin. **Segment II** with nine teeth longer and larger than those of segment III. **Segment III** with nine teeth, the distal third and fourth greatly reduced.

*Pedipalp*: **Coxa** with two ventral tubercles and one retrodorsal tubercle.

**Trochanter** with small sparse dorsal tubercles and two ventro-apical tubercles. **Femur** elongated, with small sparse dorsal setae and a ventral row of three small sparse setae; with two small setae on the ventroproximal margin. **Patella** with small sparse dorsal setae; with small prolateral, retrolateral and ventral setae on distal margin; with one small prolateral tubercle and one retrolateral small tubercle on proximal margin. **Tibia** with small sparse setae; tibial setation: prolateral IiIi and retrolateral IIi. **Tarsus** with small sparse setae; tarsal setation: prolateral IiIiiiiii and retrolateral IIiiii.

*Legs* (Figs 6D–E show leg IV): **Coxae I–III** each with one prodorsal and one retrodorsal tubercle. **Coxa IV** with sparse tubercles of different sizes and a retroventral cluster of four tubercles; with one long proapical apophysis. **Proapical apophysis of coxa IV** with sparse setae and one retroventral proximal tubercle. **Trochanters I–III** ventrally tuberculate. **Trochanters I–II** longer than wide, each with one prodorsal and one retroventral tubercle on both proximal and distal margins. **Trochanter III** as wide as long, with three prodorsal and three retroventral tubercles on distal margin. **Trochanter IV** wider than long and sparsely tuberculate; prolaterally with one conspicuous blunt central tubercle and small distal tubercles; retroventrally with one conspicuous apical tubercle and small distal tubercles. **Femora I–IV** with approximately six longitudinal rows (prodorsal, retrodorsal, retrolateral, retroventral, proventral and prolateral). **Femora I–II** straight, unarmed, with small sparse tubercles. **Femur II** with ventral tubercles near the distal margin subtly longer. **Femur III** with sparse conical tubercles of different sizes and one ventral row of tubercles increasing in size distally. **Femur IV** prolaterally curved; proximally with one dorsal cluster of 28 small tubercles and small sparse retrolateral and ventral tubercles; distally with a dorsal cluster of nine small tubercles curved posteriorly; prodorsal row of tubercles decreasing in size distally (the three basal tubercles longer); retrodorsal, retroventral, proventral and prolateral rows with small tubercles; retrolateral row with tubercles of different sizes, almost interspersed. **Dorso–basal apophysis of femur IV** inclined posteriorly, with the apex curved anteriorly; twice longer than the longest prodorsal tubercle near it and with small tubercles on the anterior face; with one small basal tubercle emerging posteriorly. **Patellae I–IV** unarmed, with small sparse tubercles. **Patellae III–IV** with the ventral tubercles curved posteriorly. **Tibiae I–IV** unarmed. **Tibiae I–II** smooth. **Tibiae III–IV** with two ventral rows of tubercles increasing in size distally, all curved posteriorly. **Tibia III** shorter than the other tibiae. **Tarsal formula**: 6(3), 12(3), 7, 8.

*Coloration in ethanol* (voucher specimen MZSP 76388): **Dorsal scutum, free tergites I–III, dorsum of coxae I–III, coxa IV, trochanters I–III, trochanter IV, femora I–IV, and patellae I–II** strong brown (centroid 55). **Tubercles of the carapace area behind the ocularium** dark olive (centroid 108). **Tubercles of areas I–III, tubercles of lateral margin of opisthosoma, posterior margin of dorsal scutum, tubercles of free tergites I–III, tubercles of coxa IV, sternites I–V, and anal operculum** brownish black (centroid 65). **Venter of coxae I–IV, genital operculum, and stigmatic area** deep orange (centroid 51). **Chelicerae, pedipalps, tibiae I–II, metatarsi I–III, third part of metatarsus IV, tarsi I–IV, anterior margin of carapace, tubercles of frontal hump, tubercles of ocularium, tubercles of lateral margin of prosoma, and ventral tubercles** brilliant greenish yellow (centroid 98). **Patellae III–IV, tibiae III–IV, and first and second parts of metatarsus IV** brownish orange (centroid 54). This coloration pattern is also observed in living specimens.

*Genitalia* (voucher specimen MZSP 76362; Figs 11D–F): **Glans** flattened and wrinkled. **Stylus** elongated and curved, with the apex dorsally projected and with subapical ventral trichomes. **Ventral process** half the stylus length and ventrally concave, with various trichomes on the margin and on the dorsal face. **Ventral plate** subrectangular and with a U-shaped apex; with three pairs of macrosetae A, one pair of macrosetae B, three pairs of macrosetae C, one pair of macrosetae D and two pairs of macrosetae E.

*Variation in males* (n = 25): **Outline of dorsal scutum** *γ*-type. **Dorsal scutum** length and width measurements variable. **Free tergites II–III** unarmed. **Prolateral apical apophysis of coxa IV** greatly reduced in length. **Femur IV** straight and with armature greatly reduced in length. **Dorso-basal apophysis of femur IV** greatly reduced in length. **Dorsal scutum coloration** from deep reddish orange (centroid 36) to strong orange yellow (centroid 68). **Leg coloration** from deep reddish orange (centroid 36) to strong orange yellow (centroid 68).

Redescription of the female (*Ilhania robusta* female paralectotype, MNRJ 42289).

*Measurements* : **DSL**: 10.0 mm. **DSW**: 7.0 mm. **LPA**: 1.0 mm.

*Dorsum* (Figs 7A and C): **Outline of dorsal scutum** *γ*-type. **Dorsal scutum** longer than wide. **Frontal hump** with two paramedian tubercles. **Area III** with two long paramedian conical spines encompassed by various tubercles.

*Venter* (Fig 7B): **Coxa I** with a proximal cluster of three prominent tubercles and a distal cluster of two prominent tubercles. **Coxa II** tuberculate, with a proximal cluster of three prominent tubercles and one distal isolated prominent tubercle.

*Chelicera*: **Fourth distal tooth of segment III** largest.

*Pedipalp*: **Femur** smooth. **Tibial setation**: prolateral Ii(Ii) and retrolateral IIi.

*Legs* (Figs 7D–E show leg IV): **Coxa III** smooth. **Coxa IV** shorter than in males; proapical apophysis absent. **Trochanter IV** without a prolateral apophysis; retroventral tubercle and prolateral proximal cluster of small setae both absent. **Femora II–III** tuberculate. **Femur IV** straight, with blunt tubercles decreasing in size distally, all curved posteriorly; dorso-basal apophysis absent. **Tibiae III–IV** with various tubercles curved posteriorly. **Tarsal formula**: 3(3), 7(3), 7, 7.

Variation in females not detected.

### Neosadocus maximus (Giltay, 1928)

Figs 1E–F, 3C, 8, 9, 11G–I, S2 Fig

*Neomitobates maximus* Giltay, 1928: 84 [55]; Kury, 2003: 135 (complete citation listing until 2003) [2]. **Type** ISNB 35, one female holotype.

*Mitoperna maxima*: Roewer, 1931: 225, fig 5 [56]; Kury, 2003: 135 (complete citation listing until 2003) [2].

*Neosadocus maximus* : Kury, 1995: 205 [15]; 2003: 135 [2]; Osses et al., 2008: 519, 520, 522, figs 1–2, 523, figs 4–5, 524, 525 [9]; Willemart et al., 2009: 52, 53, 54, figs 1–2, 55, figs 3–4, 56, fig 5, 57, 58 [10]; Rocha et al., 2011: 658, 659, 660, fig 4, 661, figs 5–5b [65]; Resende et al., 2012: 101, 102 [6]; Chelini & Machado, 2012: 1620, fig 1, 1621, 1622, 1623, fig 2, 1624, fig 3, 1625, 1626 [12]; 2014: 1148, 1149, 1150, fig 1, 1151, fig 2, 1152 [66]; Buzatto & Machado, 2014: 4 [67]; Buzatto et al., 2014: 1681 [68]; Pinto-da-Rocha et al., 2014: 522, 534 [42]; Benedetti & Pinto-da-Rocha, 2019: 462, 466, 467, 468, 469, 470 [54].

*Geraecormobius armatus* : Mello-Leitão, 1923: 137 [50]; Kury, 2003: 135 [2].

*Bunoweyhia minor* Mello-Leitão, 1935a: 19, figs 12–12a [57]; Kury, 2003: 134 (complete citation listing until 2003) [2]. Type MNRJ 41806, one male and one female syntypes. **NEW SYNONYMY**.

*Neosadocus minor* : Soares, 1945: 361 [62]; Kury, 2003: 134 (complete citation listing until 2003) [2].

**Type material.** *Neomitobates maximus* Giltay, 1928 (ISNB, one female holotype). Locality: Brazil, São Paulo, Cubatão: Piaçaguera (EXAMINED). *Bunoweyhia minor* Mello-Leitão, 1935 (MNRJ 41806, one male and one female syntypes). Locality: Brazil, São Paulo, Alto da Serra (EXAMINED).

**Other material examined.** BRAZIL. <u>São Paulo</u>: Bertioga (Vale do rio Itapanhaú), 2M 1F (MZSP 22877); Cotia (Reserva Florestal Morro Grande, Quilombo), 1M 2F (MZSP 25961); Picinguaba, 2F (MZSP 19135); Salesópolis (Estação Biológica de Boraćeia), 1M (MZSP 14344); ibidem, 1F (MZSP 19134); ibidem, 2M 1F (MZSP 16820); ibidem, 2M 1F (MZSP 17142); ibidem, 1F (MZSP 18070); ibidem, 1M (MZSP 21624); ibidem, 1F (MZSP 23628); ibidem, 4M (MZSP 23630); ibidem, 2M 2F (MZSP 23631); ibidem, 1M (MZSP 23633); ibidem, 1M (MZSP 23634); ibidem, 2M 2F (MZSP 17139); Santo Andŕe (Estação Biológica de Paranapiacaba), 1M 1F (MZSP 17176); ibidem, 1F (MZSP 30121); Ubatuba (Fazenda Capricórnio, Picinguaba), 3M 2F (MZSP 16266); ibidem, 1F (MZSP 16268); Cotia (Reserva Florestal Morro Grande), 1M 1F (MZSP 27898); Cubatão (Copebrás), 2F (MZSP 29316); ibidem, 1F (MZSP 59867); Guarujá (Serra do Guararú), 1F (MZSP 76342); ibidem, 1M (MZSP 76343); ibidem, 1M (MZSP 76344); Salesópolis (Estação Biológica de Boraćeia), 1M (MZSP 116); ibidem, 3F (MZSP 29356); ibidem, 1M (MZSP 76336); ibidem, 1F (MZSP 76337); ibidem, 1F (MZSP 76338); ibidem, 1M (MZSP 76339); ibidem, 1M (MZSP 76340); ibidem, 1F (MZSP 76341); Santo Andŕe (Estação Biológica de Paranapiacaba), 1M (MZSP 59953); ibidem, 1M (MZSP 76346); ibidem, 1M (MZSP 76347); ibidem, 1M (MZSP 76348); ibidem, 1M (MZSP 76349); ibidem, 1F (MZSP 419); ibidem, 1F (MZSP 420); ibidem, 3F (MZSP 622); ibidem, 1M (MZSP 930); ibidem, 2M 2F (MZSP 1915); ibidem, 1M (MZSP 1914); Ubatuba (Casa da Farinha, Trilha do Rio Fazenda), 1M (MZSP 76345); ibidem (Fazenda Angelim), 1F (MZSP 59841); ibidem, 1M (MZSP 76335); ibidem (Fazenda Experimental), 1M 1F (MZSP 729); ibidem, 3F (MZSP 745); ibidem, 3M 1F (MZSP 754); ibidem, 2M 1F (MZSP 759); ibidem (Morro do Cuscuzeiro, Picinguaba), 2M 1F (MZSP 16267); ibidem (Ilha das Couves), 1F (MZSP 30196).

**Distribution (Fig 2).** BRAZIL: Northern coast of the state of São Paulo. There are questionable literature records of *N. maximus* in the Southeastern region of the states of Rio de Janeiro and São Paulo. *Neosadocus* specimens were never found in any of the various subsequent Opiliones collections conducted in the Southeastern region of Rio de Janeiro state. Moreover, according to our study and as suggested by Kury (1995) [16], only *N. bufo* and *N. robustus* occur in the Southeastern region of the state of São Paulo (the latter species also occurs in the coastal region of Paraná state). Given the morphological similarities between *N. bufo* and *N. maximus*, the questionable records of the latter species may refer, instead, to records of *N. bufo*.

**Diagnosis.** *Neosadocus maximus* males resemble *N. bufo* males due to the dorsal scutum coloration pattern and the two rounded paramedian elevations densely tuberculate on area III. However, *N. maximus* males can be distinguished by the following combination of characters: short dorso-basal apophysis, same size of the longest prodorsal tubercle near it; tubercles of frontal hump, tubercles of ocularium, tubercles of carapace area behind the ocularium, tubercles of lateral margin of prosoma, and the tubercles of opisthosoma near area I pale greenish yellow (centroid 104). *N. maximus* females differ from the other *Neosadocus* females by presenting three small lateral tubercles on both sides of the anterior margin of carapace. Also, only *N. maximus* females have a small proapical apophysis on coxa IV and five long conical basal tubercles on the prodorsal row of tubercles of femur IV.

### Redescription of the male (male voucher specimen, MZSP 76344)

**Remark:** Due to the deteriorated condition of the male syntype (*Bunoweyhia minor*, MNRJ 41806), the following description is based on a specimen collected in 2013.

Photographs of the male syntype are in S2 Fig.

*Measurements* : **DSL**: 8.7 mm. **DSW**: 8.5 mm. **LDA**: 0.7 mm.

*Dorsum* (Figs 3C, 8A and C): **Outline of dorsal scutum** *γ*T-type. **Dorsal scutum** longer than wide. **Anterior margin of carapace** with three small tubercles on each side. **Ocularium** tuberculate, with two conspicuous paramedian tubercles. Carapace area behind the ocularium **tuberculate.** Lateral margin of prosoma with tubercles concentrated on the posterior half. **Areas I–II** tuberculate, each with two prominent blunt paramedian tubercles surrounded by a circle of smaller tubercles. **Area III** tuberculate, with two rounded paramedian elevations densely tuberculate, each with one central tubercle. **Lateral margin of dorsal scutum** tuberculate, with an organized external row of tubercles. **Free tergites I–III** unarmed and with tubercles of different sizes.

*Venter* (Fig 8B): **Coxa I** sparsely tuberculate; with a central longitudinal row of three prominent conical tubercles and one isolated prominent central conical tubercle; with a cluster of three prominent tubercles and one isolated prominent tubercle on distal margin. **Coxa II** sparsely tuberculate; with a cluster of three prominent tubercles and one isolated prominent tubercle on distal margin. **Coxa III** with small sparse tubercles, with an anterior row of five prominent flattened tubercles and a posterior row of ten prominent flattened tubercles, the latter almost fused with coxa IV. **Coxa IV** tuberculate. **Stigmatic area** sparsely tuberculate, with a posterior row of small tubercles. **Genital operculum** tuberculate. **Sternites I–IV** with a row of small tubercles. **Sternite V and anal operculum** tuberculate.

*Chelicera*: **Segment I** with one small tubercle; distally with one small retrolateral projection and one small prolateral projection. **Bulla** tuberculate; distally with a retrolateral cluster of two small tubercles and three small ventral tubercles. **Segment II** with seven teeth, the distal first, second and third longest. **Segment III** with eight teeth, the distal first, second and fourth longest.

*Pedipalp*: **Coxa** with one retrodorsal tubercle. **Trochanter** with two ventro-apical tubercles. **Femur** elongated, with two small dorsal setae and one small retrolateral seta. **Patella** with one small dorsal seta; with three small ventral, prolateral and retrolateral setae on distal margin; with two small ventral tubercles on proximal margin. **Tibia** with small sparse setae; prolateral IiIi and retrolateral IiIi. **Tarsus** with small sparse setae; prolateral IiIiiiii and retrolateral IIiii.

*Legs* (Figs 8D–E show leg IV): **Coxae I–II** with one prodorsal and one retrodorsal spiniform tubercle. **Coxa III** tuberculate. **Coxa IV** with sparse tubercles of different sizes and one long proapical apophysis. **Prolateral apical apophysis of coxa IV** with small dorsal tubercles and with sparse ventral setae; with one retroventral proximal tubercle and three distal inflated ventral projections. **Trochanters I–III** tuberculate, each with one prodorsal and one retroventral tubercle on distal margin. **Trochanters I–II longer** than wide, with one prodorsal and one retroventral tubercle on proximal margin. **Trochanter IV** wider than long; prolaterally with one conspicuous blunt central tubercle and small prolateral rounded tubercles on distal margin; retroventrally with one conspicuous apical tubercle and one small distal tubercle; proximally with small sparse tubercles and a prolateral cluster of five small setae. **Femora I–IV** with approximately six longitudinal rows (prodorsal, retrodorsal, retrolateral, retroventral, proventral and prolateral). **Femora I–II** straight, unarmed, with small sparse tubercles. **Femur III** with sparse conical tubercles of different sizes; distally with a dorsal cluster of six conical tubercles and two conical ventral tubercles, all curved posteriorly. **Femur IV** prolaterally curved; proximally with one dorsal cluster of 11 small tubercles, one ventral cluster of three tubercles and small sparse retrolateral tubercles; distally with a dorsal cluster of five small tubercles curved posteriorly; prodorsal, retrolateral and prolateral rows of tubercles decreasing in size distally (the five basal tubercles of prodorsal and retrolateral rows longer; the distalmost tubercle of prolateral row curved posteriorly); retrodorsal row with small tubercles; retroventral and proventral rows with tubercles of different sizes (the distalmost tubercle of retroventral row curved posteriorly; proventral row with tubercles almost interspersed). **Dorso-basal apophysis of femur IV** inclined posteriorly, with the apex curved anteriorly; the same size of the longest prodorsal tubercle near it and with small, rounded tubercles on the anterior face. **Patellae I–IV** unarmed, with small sparse tubercles. **Patellae III–IV** with the ventral tubercles curved posteriorly. **Tibiae I–IV** unarmed. **Tibiae I–II** smooth. **Tibiae III–IV** with tubercles curved posteriorly. **Tibia III** shorter than the other tibiae and with two ventral rows of tubercles increasing in size distally. **Tibia IV** with one ventral row of tubercles increasing in size distally. **Tarsal formula**: 6(3), 12(3), 7, 8.

*Coloration in ethanol* : **Ocularium, carapace area behind the ocularium, areas I–III, lateral margin of dorsal scutum, trochanters I–III, venter of coxae I–III, coxa IV, tubercles of area I and coxa IV, genital operculum, stigmatic area, and ventral tubercles** strong orange yellow (centroid 68). Anterior margin of carapace, carapace area between ocularium and lateral margin of prosoma, dorsum of coxae I–III, proapical apophysis of coxa IV, dorsum of trochanter IV, tubercles of areas II–III, tubercles of opisthosoma near areas II–III, sternites I–V, and anal operculum **brownish black (centroid 65).** Chelicerae, pedipalps, femur I, patellae I–II, tibiae I–II, metatarsi I–IV, and tarsi I–IV **brilliant greenish yellow (centroid 98).** Posterior margin of dorsal scutum, free tergites I–III, femora II–IV, patellae III–IV, tibiae III–IV, and venter of trochanter IV **brownish orange (centroid 54). Tubercles of frontal hump, tubercles of ocularium, tubercles of the area behind the ocularium, tubercles of lateral margin of prosoma, and tubercles of opisthosoma near area I** pale greenish yellow (centroid 104). This coloration pattern is also observed in living specimens.

*Genitalia* (voucher specimen MZSP 76336; Figs 11G–I): **Glans** flattened and wrinkled. **Stylus** elongated and curved, with the apex dorsally projected and with subapical ventral trichomes. **Ventral process** half the stylus length and ventrally concave, with various trichomes on the margin and on the dorsal face. **Ventral plate** subrectangular and with a U-shaped apex; with three pairs of macrosetae A, one pair of macrosetae B, three pairs of macrosetae C, one pair of macrosetae D and two pairs of macrosetae E.

*Variation in males* (n = 8): **Dorsal scutum** wider than long. **Femur IV** with four long tubercles near the dorso-basal apophysis. **Dorsal scutum coloration** brownish orange (centroid 54). **Leg coloration** brownish black (centroid 65) with patches strong orange (centroid 50).

Redescription of the female (*Bunoweyhia minor* female syntype, MNRJ 41806).

*Measurements* : **DSL**: 9.0 mm. **DSW**: 9.0 mm. **LPA**: 1.0 mm.

*Dorsum* (Figs 9A and C): **Outline of dorsal scutum** *γ*-type. **Dorsal scutum** as wide as long. **Area III** with two long paramedian conical spines surrounded by various tubercles. **Free tergites II–III** each with one central spine.

*Venter* (Fig 9B): **Coxa I** sparsely tuberculate, with a proximal cluster of three prominent tubercles and a distal cluster of two prominent tubercles on distal margin. **Stigmatic area** without a posterior row of small tubercles.

*Pedipalp*: **Femur** smooth. **Tibial setation**: prolateral Ii(Ii) and retrolateral IiIi.

**Tarsal setation**: prolateral IiIiiiii and retrolateral IIiii.

*Legs* (Figs 9D–E show leg IV): **Coxa III** smooth. **Coxa IV** shorter than in males; proapical apophysis with two small tubercles on the anterior face. **Trochanter IV** without a prolateral apophysis, retroventral tubercle and prolateral proximal cluster of small setae both absent. **Femora II–III** tuberculate. **Femur IV** straight, with conical tubercles decreasing in size distally, all curved posteriorly; dorso-basal apophysis absent. **Tibiae III–IV** with various tubercles curved posteriorly. **Tarsal formula**: 3(3), 7(3), 7, 8.

Variation in females not detected.

### *Neosadocus misandrus* (Mello-Leitão, 1934)

**Fig 10**

*Metagonyleptes misandrus* Mello-Leitão, 1934: 416, fig 7 [69]; Kury, 2003: 135 (complete citation listing until 2003) [2]; Coronato-Ribeiro et al., 2013: 505, 513, fig 5 [53]. **Type**

IBSP 12, one female holotype.

*Neosadocus misandrus* : Kury, 2003: 135 [2].

**Type material.** *Metagonyleptes misandrus* Mello-Leitão, 1934 (IBSP 12, one female holotype). Locality: Brazil, Unknown state (EXAMINED).

**Distribution.** BRAZIL: There is not a specific locality for the holotype.

**Diagnosis.** *Neosadocus misandrus* is only known from the female holotype, which can be distinguished from the other females of the genus by the following characters: (i) two long paramedian conical spines on the ocularium (Figs 10A–B), whereas all other females have two conspicuous paramedian tubercles on the ocularium; (ii) absence of a central spine on free tergites II and III, whereas they are present in all other females; and (iii) small blunt tubercles of the same size on femur IV, all curved posteriorly (Figs 10C–D), whereas the tubercles of all other females decrease in size distally and may be blunt (*N. bufo* and *N. robustus*) or conical (*N. maximus*).

### Molecular data, phylogenetic analyses and divergence times

Our dataset comprises 83 COI sequences (42 haplotypes) and 96 ITS2 sequences (32 haplotypes) with 505 base pairs (bp) and 373 bp (with gaps), respectively (Table 1, S2 Table). No stop codons or ambiguous peaks were detected in the COI dataset, suggesting the absence of pseudogenes or numts in the sequences.

The best-fit substitution models for COI and ITS2 datasets were HKY+I+G and HKY+I, respectively. Both Maximum Likelihood and Bayesian phylogenetic analyses revealed gene trees with similar topologies (Fig 12; topologies recovered in ML analyses are shown in S3 Fig). The COI cladogram supported the monophyly of *Neosadocus*, showing *N. bufo*, *N. maximus* and *N. robustus* as highly supported distinct clades; in the ITS2 tree, however, two outgroup species (*Iguapeia melanocephala* and *Iporangaia pustulosa*) were positioned within the *Neosadocus* clade (which may be a result of the close relationship among these species and lower divergence observed in the nuclear sequences).

The multilocus calibrated tree recovered the monophyly of *Neosadocus*, corroborated its close relationship with *Iguapeia melanocephala* and *Iporangaia pustulosa*, and indicate that *Neosadocus* species diversified in the Miocene, with divergences starting at ca. 17.14 Mya [95% highest posterior density (HPD) = 8.88–26.94 Mya, Fig 13; calibrated COI and ITS2 trees are provided in S4 Fig and S5 Fig, respectively].

### Intraspecific analyses: population structure and genetic diversity

The COI median-joining network recovers several haplogroups separated by a great number of mutational steps in all three species, while ITS2 haplotypes are separated by only one to three nucleotide differences (Fig 14). Most COI haplotypes were exclusive (i.e., detected in one single location), except for three haplotypes found in populations geographically close (H25, in Morretes and Paranaguá; H26, in Antonina and Morretes; and H32, in Cajati and Barra do Turvo; S2 Table). For ITS2, seven out of the 32 haplotypes were shared among different locations (H2, H7, H17, H19, H20, H21 and H30; in all cases, the populations were geographically close).

**Fig 14.**
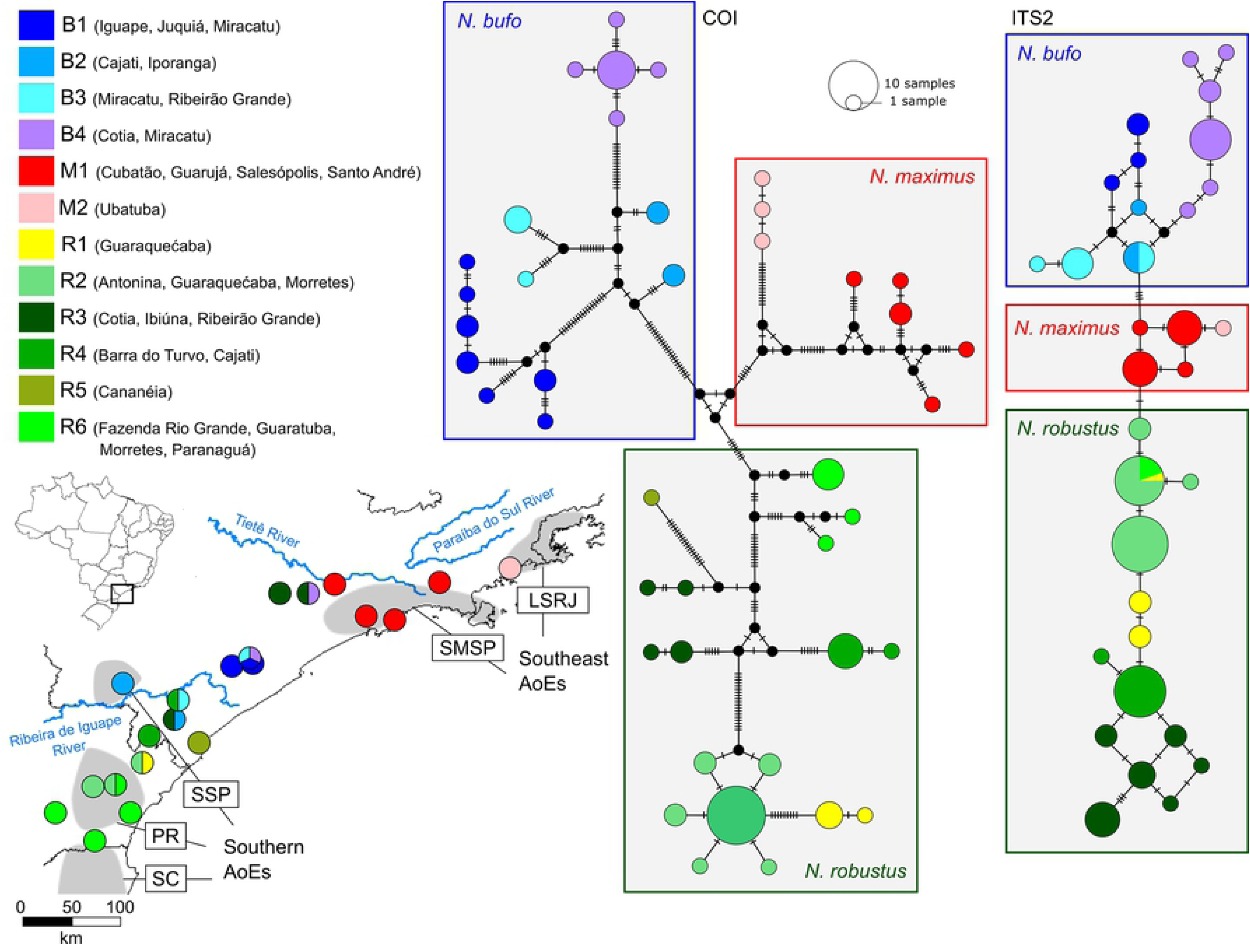
Haplotype networks and structure obtained with BAPS. The colors represent the genetic clusters indicated for COI sequences: *N. bufo* = B1–B4; *N. maximus* = M1–M2; *N. robustus* = R1–R6 (the information was extrapolated for ITS2 data, which exhibited a less marked subdivision). In the map, the main rivers and Areas of Endemism (AoEs) mentioned in the discussion are shown (LSRJ = Southern Rio de Janeiro Coast; SMSP = Serra do Mar of São Paulo; SSP = Southern São Paulo; PR = Parańa; SC = Santa Catarina; adapted from [20]).

The BAPS results indicated four COI clusters for *N. bufo* (groups B1–B4), two for *N. maximus* (M1 and M2), and six for *N. robustus* (R1–R6; Fig 14, S2 Table), which matched the main lineages detected in the phylogenetic analyses (Fig 12). These intraspecific clusters are spatially distinct, although they overlap in few regions (as in Miracatu, Guaraqueçaba and Morretes, populations composed by sequences from different BAPS clusters; Fig 14). Two *N. bufo* clusters are distributed in the Northern portion of the species range (B1, B4), while the other two are in the Southern region (B2, B3). In the case of *N. maximus*, samples from the Ubatuba population (at the Northern edge of the species distribution) formed a separate cluster; this result must be treated with caution, since there is an indication of isolation-by-distance in this species (see below). Finally, *N. robustus* clusters were predominantly distributed to the North (R3, R4, R5) or South (R1, R2, R6) of the Ribeira do Iguape River region and the Northern clusters sympatric with *N. bufo* (Fig 14). The structure pointed by BAPS for ITS2 sequences was similar but less marked (which is probably due to the higher similarity among sequences), showing the same subdivision for *N. maximus*, three groups for *N. bufo* (merging B2+B3 COI clusters), and three for *N. robustus* (the first combining R1+R2+R6 COI clusters and the other two similar to R3 and R4, except for some samples from Ribeirão Grande and Ibiúna; S2 Table). Given that there was more structure in the COI sequences than in the ITS2 (S2 Table), we extrapolated the geographical distribution of the mitochondrial clusters in the ITS2 haplotype network shown in Fig 14.

The genetic distances among samples were considerably greater for the mitochondrial datasets. The mean distances between species varied from 39.59 to 46.91 average pairwise differences for COI sequences, and from 4.02 to 8.01 differences for the ITS2 dataset (S3 Table and S4 Table). Within each species, the average number of pairwise COI differences between populations varied from 7.6 to 33.0 in *N. bufo*; from

8.0 to 33.0 in *N. maximus* ; and from 0.5 to 31.0 in *N. robustus*. For the ITS2 sequences, the greatest genetic distances between populations were 7.3, 3.0 and 7.7 for *N. bufo*, *N. maximus* and *N. robustus*, respectively (S5–S10 Tables). The Mantel tests indicated a significant correlation between genetic and geographic distances for the COI sequences of *N. maximus* (r2 = 0.8, *p* = 0.007), and ITS2 sequences for all species analyzed (*N. bufo*: r2 = 0.55, *p* = 0.035; *N. maximus* : r2 = 0.75, *p* = 0.049; *N. robustus* : r2 = 0.74, *p* = 0.0002).

The global *ϕ*_ST_ values were high for the three species, indicating strong population structure (*N. bufo*: *ϕ*_ST_COI = 0.76, *ϕ*_ST_ITS2 = 0.66; *N. maximus*: *ϕ*_ST_COI = 0.96, *ϕ*_ST_ITS2 = 0.87; *N. robustus* : *ϕ*_ST_COI = 0.82, *ϕ*_ST_ITS2 = 0.75). The significant pairwise *ϕ*_ST_ values varied from 0.252 to 1.0 for the COI samples, and from 0.168 to 1.0 for the ITS2. The lowest values were found for *N. robustus* samples collected between Morretes and Guaraqueçaba (S11–S16 Tables); most values, however, were statistically non-significant due to the small size of the samples.

The haplotype diversity values were high for the three species, varying from 0.88 to 0.97 for COI sequences, and from 0.74 to 0.9 for ITS2 (2). The nucleotide diversity obtained for COI sequences was substantially higher, varying from 0.034 to 0.041 in each species, while the ITS2 values varied from 0.003 to 0.01 nucleotide differences. Within municipalities, the genetic variation was generally low for both markers, since most populations presented only one or few closely related haplotypes.

## Discussion

This is the first taxonomic revision and molecular phylogenetic study focusing on *Neosadocus*. Herein, we discuss the different aspects of our results, from intraspecific variation in secondary sexual characters in males to species relationships and population genetic structure. We also inferred historical events in the Brazilian Atlantic Forest that possibly influenced the distribution of *Neosadocus* species and their evolutionary patterns.

### Male polymorphism in *Neosadocus robustus*

Three types of morphological polymorphism are recognized among gonyleptid harvestmen: sexual dimorphism, male polymorphism and variation in non-sexual/non-fixed traits [70]. All *Neosadocus* species present sexual dimorphism. However, only *N. robustus* show male polymorphism (S1 Fig), a phenomenon that had been previously documented for other harvestmen but not for this species [67, 68, 70–72]. We observed morphological polymorphism in the following traits of male specimens of *N. robustus* : outline, length and width of dorsal scutum (*γ*T-type in major males; *γ*T-type in minor males); presence or absence of a central spine on free tergites II and III; length of prolateral apical apophysis of coxa IV (long in major males; reduced in minor males); curvature and armature of femur IV (curved and strongly armored in major males; straight and with reduced armature in minor males); coloration of dorsal scutum and legs. Willemart et al. (2009) [10] reported that the armature of the fourth pair of legs is used by *Neosadocus maximus* males to fight for resources and nesting sites defense [51] -. We hypothesize that this phenomenon occurs in major males of *N. robustus*. Nevertheless, the sneaky behavior had been previously documented in minor males of the harvestmen *Serracutisoma proximum* [73]: minor males invade the territories of major males and furtively mate with egg-guarding females. Guarding eggs had been previously documented for females of *N. maximus* [11, 12]. If *N. robustus* females also guard their eggs, we conjecture that they also mate with minor males of *N. robustus*. These cases and hypotheses illustrate the male polymorphism associated with different reproductive strategies [74]. Finally, we also observed an apparent polymorphism of non-sexual/non-fixed traits in males: presence/absence of a central spine on free tergites II and III; coloration of dorsal scutum and legs. Coloration was the only variation we observed in both males and females. We do not believe that the presence/absence of a central spine on free tergites II and III has anything to do with fighting in males, since this trait is variable among major and minor males. We do not have a reasonable hypothesis to explain color variation.

### Molecular phylogenetics, divergence times and biogeographic inferences

The molecular analyses indicated a clear differentiation among *Neosadocus bufo*, *N. maximus* and *N. robustus* (Fig 13), corroborating the morphological evidence that these species are valid. The results of the multilocus Bayesian analysis suggests that *N. maximus* and *N. robustus* are more closely related to each other than each is to *N. bufo*; however, this result needs to be looked at with caution, since the posterior probability of this clade is relatively low (0.82), as in all other phylogenetic trees (Fig 12, S3 Fig, S4 Fig, S5 Fig). The inclusion of more molecular data and a morphological cladistic analysis may assist the elucidation of the relationships among the species of *Neosadocus*. Our results also show a close relationship between *Neosadocus* and other Progonyleptoidellinae species, especially *Iguapeia melanocephala* and *Iporangaia pustulosa*. This proximity had been previously hypothesized by Pinto-da-Rocha et al. (2014) [42] and Benedetti & Pinto-da-Rocha (2019) [54], also based on molecular data. Caetano & Machado (2013) [75] also suggested that *Neosadocus* and Progonyleptoidellinae are closely related, based on behavioral, chemical and ecological data. Since this work aimed to uncover the phylogenetic relationships of *Neosadocus* species, we will not discuss in detail the phylogenetic relationship between *Neosadocus* and Progonyleptoidellinae. More data are still necessary to further resolve that aspect of the phylogeny.

Most of the genetic divergence in *Neosadocus* occurred in the Miocene [speciation events at ca. 12–17 Mya (95% HPD = 6–27 Mya); main intraspecific divergences starting at ca. 7–14 Mya (95% HPD = 4–25 Mya; Fig 13, S2 Fig, S3 Fig). This period was marked by intense orogenic activity in the Southern Atlantic Forest, which may have driven the evolutionary history of *Neosadocus*. An important geological structure in the region is known as the Continental Rift of Southeastern Brazil, which includes important present-day relief components, as the Ribeira do Iguape and Paráıba do Sul River Valleys, and the Serra do Mar and Serra da Mantiqueira mountain ranges. Neogene-Quaternary geomorphologic processes along this rift probably increased the topographical differences between mountains and valleys, rearranging rivers and lake systems [76, 77]. Other historical events rather than tectonism, for instance changes in climatic conditions (partially imposed by the above-mentioned relief modifications) and marine transgressions [78] may also have played a role in the differentiation of *Neosadocus* populations, since they could have influenced forest rearrangements or reduction to isolated patches. A phylogeographic study with other Southern Atlantic Forest harvestmen (A*cutisoma longipes*) corroborates that the drier conditions in valleys (in comparison to Serra do Mar mountain slopes) may have limited the gene flow of populations [22], a consequence of the considerable dependence on humidity and poor dispersal ability of these organisms, promoting lineage diversification.

Studies on the historical relationships among the AoEs of harvestmen in the Atlantic Forest have proposed not only that all the processes mentioned above might be involved in the regionalization of the opiliofauna, but also that they acted as reiterative barriers, which is reflected in the current complex biogeographic patterns with spatial congruence in some areas [7, 20]. Using cladistic biogeographic methods, DaSilva et al. (2017) [20] detected an important split between the AoEs mostly at the north and south of Ribeira de Iguape River Valley region (named Southeastern and Southern blocks, respectively). The current distributions of *Neosadocus* species corroborate this division: *N. maximus* is restricted to the Northern edge of the range of *Neosadocus* [being restricted to Serra do Mar of São Paulo (SMSP) and Southern Rio de Janeiro Coast (LSRJ) area, in the Southeastern block], while the distribution of *N. bufo* and *N. robustus* partially overlap and occupy the South-Central portion of the genus’ distribution, on the AoEs of the Southern block [Southern São Paulo (SSP), Parańa (PR) and Santa Catarina (SC), although many individuals were sampled out of AoEs’ congruence cores; Fig 14]. This geographical delimitation, associated with the old (Miocene) divergence times estimated, suggests that ancient historical barriers have influenced the diversification of harvestmen in this region.

Several phylogeographic breaks in the Southern Atlantic Forest had been previously reported for different taxa. A variety of groups exhibit genetic disjunctions that are geographically similar to *Neosadocus* around the Continental Rift of Southeastern Brazil, as birds [79–81], mammals [82], amphibians [83–86], snakes [87], insects [88, 89], and planarians [90]. However, the breaks observed in this study are considerably older than the divergences reported for most taxa, a pattern already observed for other co-distributed harvestmen [21–23]. This time-scale heterogeneity reveals that the evolution of the Atlantic Forest biota is complex and cannot be attributed to a single process; rather, it is more reasonable that multiple historical barriers arose reiteratively in a same region. Additionally, spatial and temporal incongruences may result from the different ways species respond to environmental changes, responses that are mediated by taxon-specific traits, dispersal ability, population dynamics, physiological constraints, and/or ecological interactions [91]. Hence, organisms with extremely low vagility, philopatry and humidity-dependence, as harvestmen, tend to retain relatively older phylogeographic signals with respect to other groups, which reinforces the value of these species for comparative biogeographical studies. As different co-distributed taxa are more broadly investigated, it becomes clear that both Quaternary events (as the global climatic oscillations that promoted cycles of forests shifts) and older processes (as the Neogene geomorphological and climatic episodes described above) acted in combination to produce taxon-specific idiosyncratic patterns of diversification in the Atlantic Forest [92].

### Population structure and genetic diversity

All three species exhibited great variability and strong population structure. Most haplotypes are geographically restricted to a single location, and the few exceptions are shared among close locations (Fig 14), a result that matches the high genetic distances and *φ*_ST_ values. This pattern is similar to those reported for other Atlantic Forest harvestmen [21–23]. Tropical harvestmen, with the exception of some members of the suborder Eupnoi [93], have low vagility - a rare case among arthropods of similar size [3, 5, 6]. Moreover, they are very humidity-dependent and only survive within a narrow temperature zone [5]. All these characteristics limit their chances of dispersal and contribute to the geographical structure commonly found in their populations. The restricted opportunities for relocation may be more intense in regions with valleys (as the Southern Atlantic Forest), since these lower altitude areas are consistently more unstable and are usually drier than the forests inhabited by these harvestmen.

Our BAPS results indicated genetic subdivisions within each species, especially as calculated using COI sequences. Although the genetic groups overlap somewhat, they are geographically separated (Fig 14), which suggests that historical events might have reduced (or interrupted) the gene flow among some populations in the past (as discussed above). This intraspecific differentiation is consistent with the AoEs proposed for harvestmen, corroborating the biotic disjunctions already detected [7, 20]: the M1 and M2 groups of *N. maximus*, for example, occur in SMSP and LSRJ areas, respectively; similarly, the intraspecific groups of *N. bufo* and *N. robustus* are spatially congruent with different Southern AoEs blocks (although the association for these two species is less clear; Fig 14). However, as we obtained statistically significant results in Mantel tests for some datasets (especially for ITS2 sequences), we cannot completely discard that these intraspecific groups reflect isolation-by-distance rather than historical barriers.

Although our morphological observations indicate greater variation in *N. robustus*, the total genetic diversity was similar among the three species (Table 2). Also, we found no correlation between the morphological variation and the genetic groups pointed by BAPS in any species, indicating that such variability is not the result of restricted gene flow. More studies correlating morphological traits and a greater amount of neutral and non-neutral molecular markers could bring more robust results on this issue.

**Table 2.** Haplotype and nucleotide diversities for each *Neosadocus* species (gray lines) and per location (under species indices). N = number of specimens; H = number of haplotypes; S = number of segregating sites; Hd (s.d.) = haplotype diversity (standard deviation); ***π*** (s.d.) = nucleotide diversity (standard deviation).

| Species/<br>Municipality | COI |  |  |  |  | ITS2 |  |  |  |  |
| --- | --- | --- | --- | --- | --- | --- | --- | --- | --- | --- |
| | N | H | S | HD (s.d.) | $\pi$ (s.d.) | N | H | S | HD (s.d.) | $\pi$ (s.d.) |
| ***Neosadocus bufo*** | **30** | **16** | **67** | **0.931 (0.028)** | **0.041 (0.021)** | **27** | **13** | **12** | **0.9 (0.037)** | **0.01 (0.006)** |
| Cajati | 2 | 1 | 0 | - | - | 1 | 1 | 0 | - | - |
| Cotia | 2 | 2 | 4 | 1 (0.5) | 0.008 (0.009) | 2 | 2 | 2 | 1 (0.5) | 0.005 (0.007) |
| Iguape | 8 | 5 | 12 | 0.893 (0.086) | 0.01 (0.006) | 3 | 2 | 1 | 0.667 (0.314) | 0.002 (0.002) |
| Iporanga | 2 | 1 | 0 | - | - | 2 | 1 | 0 | - | - |
| Juquiá | 1 | 1 | 0 | - | - | - | - | - | - | - |
| Miracatu | 10 | 5 | 46 | 0.667 (0.163) | 0.022 (0.012) | 12 | 6 | 9 | 0.682 (0.148) | 0.005 (0.004) |
| Ribeirão Grande | 5 | 1 | 0 | - | - | 7 | 3 | 3 | 0.667 (0.16) | 0.003 (0.003) |
| ***Neosadocus maximus*** | **9** | **8** | **45** | **0.972 (0.064)** | **0.038 (0.021)** | **13** | **5** | **3** | **0.744 (0.087)** | **0.003 (0.003)** |
| Cubatão | 1 | 1 | 0 | - | - | 1 | 1 | 0 | - | - |
| Guarujá | 3 | 2 | 1 | 0.667 (0.314) | 0.001 (0.002) | 1 | 1 | 0 | - | - |
| Salesópolis | 1 | 1 | 0 | - | - | 6 | 2 | 1 | 0.333 (0.215) | 0.001 (0.001) |
| Santo André | 1 | 1 | 0 | - | - | 4 | 1 | 0 | - | - |
| Ubatuba | 3 | 3 | 2 | 1 (0.272) | 0.003 (0.003) | 1 | 1 | 0 | - | - |
| ***Neosadocus robustus*** | **44** | **18** | **67** | **0.88 (0.039)** | **0.034 (0.017)** | **56** | **14** | **9** | **0.873 (0.023)** | **0.008 (0.005)** |
| Antonina | 7 | 1 | 0 | - | - | 8 | 2 | 2 | 0.429 (0.169) | 0.002 (0.002) |
| Barra do Turvo | 2 | 1 | 0 | - | - | 6 | 1 | 0 | - | - |
| Cajati | 4 | 2 | 2 | 0.5 (0.265) | 0.002 (0.002) | 6 | 2 | 1 | 0.333 (0.215) | 0.001 (0.001) |
| Cananéia | 1 | 1 | 0 | - | - | - | - | - | - | - |
| Cotia | 1 | 1 | 0 | - | - | 4 | 1 | 0 | - | - |
| Fazenda Rio Grande | 1 | 1 | 0 | - | - | 1 | 1 | 0 | - | - |
| Guaraqueçaba | 8 | 5 | 14 | 0.857 (0.108) | 0.014 (0.008) | 9 | 4 | 4 | 0.778 (0.11) | 0.005 (0.003) |
| Guaratuba | 1 | 1 | 0 | - | - | - | - | - | - | - |
| Ibiúna | 1 | 1 | 0 | - | - | 2 | 1 | 0 | - | - |
| Morretes | 12 | 4 | 29 | 0.652 (0.133) | 0.01 (0.006) | 10 | 2 | 1 | 0.467 (0.132) | 0.001 (0.001) |
| Paranaguá | 3 | 1 | 0 | - | - | 2 | 1 | 0 | - | - |
| Ribeirão Grande | 3 | 2 | 1 | 0.667 (0.314) | 0.001 (0.002) | 8 | 5 | 5 | 0.857 (0.108) | 0.005 (0.003) |
| **Total** | **83** | **42** | **130** | **0.957 (0.013)** | **0.067 (0.033)** | **96** | **32** | **22** | **0.945 (0.01)** | **0.014 (0.007)** |

## Conclusion

Based on morphological observations and molecular phylogenetic analyses, our study confirmed the taxonomic validity of three *Neosadocus* species: *N. bufo*, *N. maximus* and *N. robustus*. The latter species presented a high degree of male polymorphism, mostly related to reproductive strategies. Although morphological observations corroborated the validity of *N. misandrus*, the absence of specimens besides the female holotype did not allow us to include this species in our phylogenetic analysis.

Furthermore, our time-calibrated inferences and phylogeographic analyses suggest Neogene speciation events, deep intraspecific lineages and strong population structure in all three species. These results match the patterns commonly found for Neotropical harvestmen and corroborate that historical tectonic and climatic events might have imposed reiterative or longstanding barriers to the gene flow among harvestmen in the Atlantic Forest areas.

## Supporting information

**S1 Table. Primers used for the amplification of each molecular marker.**

**S2 Table. COI and ITS2 haplotypes and BAPS results.**

**S3 Table. Genetic distances between *Neosadocus* species obtained for COI sequences.**

**S4 Table. Genetic distances between *Neosadocus* species obtained for ITS2 sequences.**

**S5–S10 Tables. Genetic distances between *Neosadocus* populations obtained for COI and ITS2 sequences.**

**S11–S16 Tables. Pairwise ΦST values between *Neosadocus* populations obtained for COI and ITS2 sequences.**

**S1 Fig. Morphological polymorphism in males of *Neosadocus robustus***. Scale bars: 1 mm.

**S2 Fig. *Neosadocus maximus* (*Bunoweyhia minor* male syntype, MNRJ 41806).** A. Dorsal view. B. Ventral view. C. Left lateral view. D–E. Right trochanter–patella IV (D: prolateral view; E: retrolateral view). Scale bars: 1 mm.

S**3 Fig. COI and ITS2 topologies recovered with RAxML.**

**S4 Fig. COI calibrated topology recovered in *BEAST analysis.**

**S5 Fig. ITS2 calibrated topology recovered in *BEAST analysis.**

## Acknowledgments

We thank Alípio R. Benedetti, André A. Nogueira and Giulia M. Ribeiro for their support during fieldwork; Beatriz V. Freire and Manuel A. Junior for their help and advice on DNA extraction, amplification and sequencing; Enio Mattos and Phillip Lenktaits for helping us take scanning electron microscope photographs. We also thank Dr. Adriano Brilhante Kury for the loan of the type material deposited on MNRJ; Dr. Antonio D. Brescovit the access to the type material deposited at IBSP; and Dr. Yves Samin for providing high quality photographs of the type material deposited at ISNB. We also thank Dr. Márcio B. da Silva, Dr. Marcos R. Hara and M.Sc. Wendy Y. Arroyo Pérez for all their remarkable observations and comments on a previous version of this manuscript.

